# Barriers to gene flow play an important role in miantaining reproductive isolation between two closely related *Populus* (Salicaceae) species

**DOI:** 10.1101/2020.09.08.287292

**Authors:** Yang Tian, Shuyu Liu, Pär K. Ingvarsson, Dandan Zhao, Li Wang, Baoerjiang Abuduhamiti, Zhiqiang Wu, Jianguo Zhang, Zhaoshan Wang

## Abstract

In most species, natural selection plays a key role in genomic heterogeneous divergence. Additionally, barriers to gene flow, such as chromosomal rearrangements or gene incompatibilities, can cause genome heterogeneity. We used genome-wide re-sequencing data from 27 *Populus alba* and 28 *P. adenopoda* individuals to explore the causes of genomic heterogeneous differentiation in these two closely related species. In highly differentiated regions, neutrality tests (Tajima’s D and Fay & Wu’s H) revealed no difference while the absolute divergence (d_xy_) were significantly higher than genome background, which indicates that natural selection did not play a major role but barriers to gene flow play an important role in generating genomic heterogeneous divergence and reproductive isolation. The two species diverged ∼5-10 million years ago (Mya), when the Qinghai-Tibet Plateau reached a certain height and the inland climate of the Asian continent became arid. We further found some genes that are related to reproduction.

## Introduction

Understanding the origins of reproductive isolation is a major goal of modern speciation studies in plants because reproductive isolation is crucial for speciation (Cruickshank and Hahn, 2014; Widmer et al., 2008). Postzygotic isolation may be caused by inferior niche adaptation, reduced attractiveness of hybrids or reduced fertility and/or viability of hybrids resulting from intrinsic incompatibilities (Eric, 2015; Levin, 1978; Lynch and Force, 2000; MacNair and Christie, 1983; Orr and Turelli, 2001; Qvarnström and Bailey, 2008). Some degree of post-zygotic isolation often occurs in recently diverged species and are followed by the slow establishment of stronger prezygotic isolation (Eric, 2015; Wang et al., 2016; Widmer et al., 2008).

The origins of reproductive isolation include multiple evolutionary forces, including genetic drift, natural selection and mutations (Eric, 2015). Reproductive isolation is often considered to be an important by-product of genetic drift or differential adaptation when no gene flow is present (Funk et al., 2016; Noor et al., 2001; Nosil et al., 2008). However, divergent selection in the face of gene flow is a common scenario that can have different effects across the genome of an organism (Nosil et al., 2009). Positive and/or negative selection at individual loci can be reduced in one or both daughter populations, thereby increasing the degree of differentiation at these loci (Geraldes et al., 2011; Rifkin et al., 2019). Selection on linked sites (due to genetic hitchhiking or background selection) can also lead to more differentiation in genomic regions with lower recombination rates (Nachman Michael and Payseur Bret, 2012). In addition, phenomena such as chromosomal rearrangements or gene incompatibilities can also contribute to reproductive isolation (Baack et al., 2015; Noor et al., 2001; Rieseberg, 2001; Schumer et al., 2014). Gene incompatibilities may lead to infertility or even mortality of hybrids between species and chromosomal rearrangements are known to play an important role in the origin of reproductive isolation because they sometimes disrupt meiosis in hybrids, resulting in inviability and/or infertility in hybrids (Hou et al., 2014; Lynch and Force, 2000; Teresa Avelar et al., 2013).

A common interpretation of the small number of strongly differentiated regions identified during genome scanning is that they represent loci related to reproductive isolation or ecological specialization and are that these loci are under strong selection to prevent introgression between species (Cruickshank and Hahn, 2014; Payseur and Rieseberg, 2016; Wu, 2001). Loci unrelated to reproductive isolation will experience homogenization from gene flow and will consequently exhibit low or no genetic differentiation. These differences in the action of natural selection is thought to explain a pattern of heterogeneous differentiation among sites in the genomes of closely related species (Cruickshank and Hahn, 2014). There are many factors leading to heterogeneous genomic divergence (Nosil et al., 2009), including selection due to ecological causes (Funk et al., 2016) or genetic conflicts (Nosil et al., 2009; Presgraves, 2007), random effects of genetic drift (Funk et al., 2016), variable mutation rates (Cruickshank and Hahn, 2014; Lynch and Force, 2000) or the genomic distribution and effect size of selected genes and chromosome structure (Noor et al., 2001; Rieseberg, 2001; Teresa Avelar et al., 2013).

Therefore, it is important to analyse different driving factors to account for how heterogeneous genomic divergence is formed and what roles they play in the formation and maintenance of reproductive isolation and adaptive phenotypic differentiation (Jacobs et al., 2018). In the absence of gene flow and natural selection between species that have recently diverged, the heterogeneous differentiation of the genome is simply resulting from stochastic variation in coalescence times (Barton, 2006). However, if selection is present, regions that experience strong natural selection will exhibit larger differences than those under weaker selection and will thus share less ancestral variation (Cruickshank and Hahn, 2014). In the presence of gene flow, selection reduces the diversity of selected loci but also on sites linked to the selected loci thereby promoting genetic differentiation and resulting in an inflation of measures such as *F*_ST_ between species (Cruickshank and Hahn, 2014; Jakobsson et al., 2013; Walsh and Blows, 2009). Natural selection reduces effective gene flow near selected sites; however, selection does not affect the redistribution of alleles in daughter species. Other regions exhibit low differentiation, either because they maintain a high level of ancestral polymorphism or because the homogenizing effects of gene flow, thereby leading to heterogenous genetic differentiation across the genome or an organism (Turner and Hahn, 2010; Wang et al., 2016).

Barriers to gene flow, such as chromosomal rearrangements or gene incompatibilities, can also cause genome heterogeneity because they inhibit the redistribution of alleles between daughter species. Such barriers can lead to the formation of highly differentiated regions by countering the homogenizing effects of gene flow (Feder and Nosil, 2009). At the same time, other areas of the genome are homogenized by gene flow, consequently showing lower levels of differentiation. This process results in a highly variable degree of genetic differentiation throughout the genome, also known as heterogeneous genomic differences (Nosil et al., 2009; Ortíz-Barrientos et al., 2002; Wolf and Ellegren, 2016). Chromosomal rearrangements are also known to reduce the fitness of heterozygotes via by affecting meiosis, which may eventually result in reproductive isolation (Baack et al., 2015).

In summary, all three factors outlined above can explain heterogeneity in genomic differentiation and distinguishing between them and deciding which are the major factors in a particular scenario is a challenging task. Of course, these assumptions are not mutually exclusive and detailed information about the process of speciation, such as the time of speciation, the time of differentiation, and the demographic history of the species pair, are required to understand the causes of heterogeneous differentiation across the genome (Nosil et al., 2008; Wang et al., 2016).

*Populus alba* and *P. adenopoda* are two important tree species in the section *Populus* with large ecological and economic value (Wang et al., 2010). A phylogenetic tree constructed using a combination of cpDNA and nuclear DNA show that the two species are sister species (Wang et al., 2014), indicating that they have diverged recently. *P. alba*, white poplar, is widely distributed and cultivated across the North African deserts, European flood plain forests and in central Asian regions having a strong continental climate and severe winter frosts (Dickmann and Kuzovkina, 2014; Stölting et al., 2015). *P. adenopoda* is endemic to China, where it often occurs on sunny slopes or along riversides and is common in the southwest, where it shows good natural regeneration (Fan et al., 2018). Natural populations of *P. alba* often hybridise with populations of other, closely related species and hybridization with *P. adenopoda*, have produced a large number of natural *P. tomentosa* hybrids, which have poor fertility and produce almost no offspring (Li et al., 1997; Wang et al., 2011). Therefore, while the two species have diverged recently they nevertheless show evidence for very strong postzygotic isolation. *P. alba* and *P. adenopoda* is thus a promising system for evaluating how different evolutionary forces contribute to the genomic patterns of divergence during speciation and how the accumulation of genetic differences leads to intrinsic barriers to reproduction (Palumbi, 1994).

In this study, we used genome-wide resequencing data from the two species to infer their divergence time and estimate their historical population dynamics. By examining population genomic statistics, such as genetic diversity and relative and absolute divergence, we expect to identify the major factors leading to heterogeneous genomic differentiation between the two species. We also identify outlier regions and genes that may be associated with adaptation to novel environments during the speciation process.

## Results

A total of 27 *P. alba* and 28 *P. adenopoda* individuals were selected for whole-genome resequencing. The *P. trichocarpa* genome (v3.0) was used as the reference genome because of its high-quality assembly and annotation as well as the high genomic synteny between members of sections *Populus* (*P. alba* and *P. adenopoda*) and *Tacamahaca* (*P. trichocarpa*) (Liu et al., 2019; Pakull et al., 2009; Tuskan et al., 2006). After removing reads containing adaptors and then trimming and filtering, the sequenced reads from the two species were mapped to the *P. trichocarpa* reference genome (v3.0) (Tuskan et al., 2006) with a moderately high mapping rate (84.3% on average, Table S1). The mean coverage of reads that were uniquely mapped per site was 23.9 and 24.8 in *P. alba* and *P. adenopoda*, respectively (Table S1). In this study, we used two complementary bioinformatics methods. First, the genetic statistics of populations that depend on the inferred site-frequency spectrum (SFS) were estimated directly from the genotype likelihoods rather than by calling the genotype (Nielsen et al., 2012), in ANGSD (Korneliussen et al., 2014). Second, HaplotypeCaller in GATK (Danecek et al., 2011) was used to call single nucleotide polymorphisms (SNPs) and genotypes for estimates that require accurate genotype calls. VCFtools (Danecek et al., 2011) was used to perform a series of filtering steps on the VCF files prior to data analysis. A total of 22,525,740 and 21,217,984 high-quality SNPs were obtained from the 27 *P. alba* and 28 *P. adenopoda* samples, respectively, after filtering.

### Population structure and Demographic histories

NGSadmix (Skotte et al., 2013) was used for quantifying admixture proportions for the two species. The program infers individual ancestry based on genotype likelihoods and examines the genetic relationships between individuals. The analyses were run with the number of genetic clusters (K) varying from 2 to 4. When K=2 (Figure 1b), all samples were divided into two species-specific populations. When K=3 (Figure 1b), the results were no longer meaningful. This result was verified by principal component analysis (PCA) that show a clear separation of the two species and little evidence for further subdivision within species (Figure 1c).

**Figure 1.**
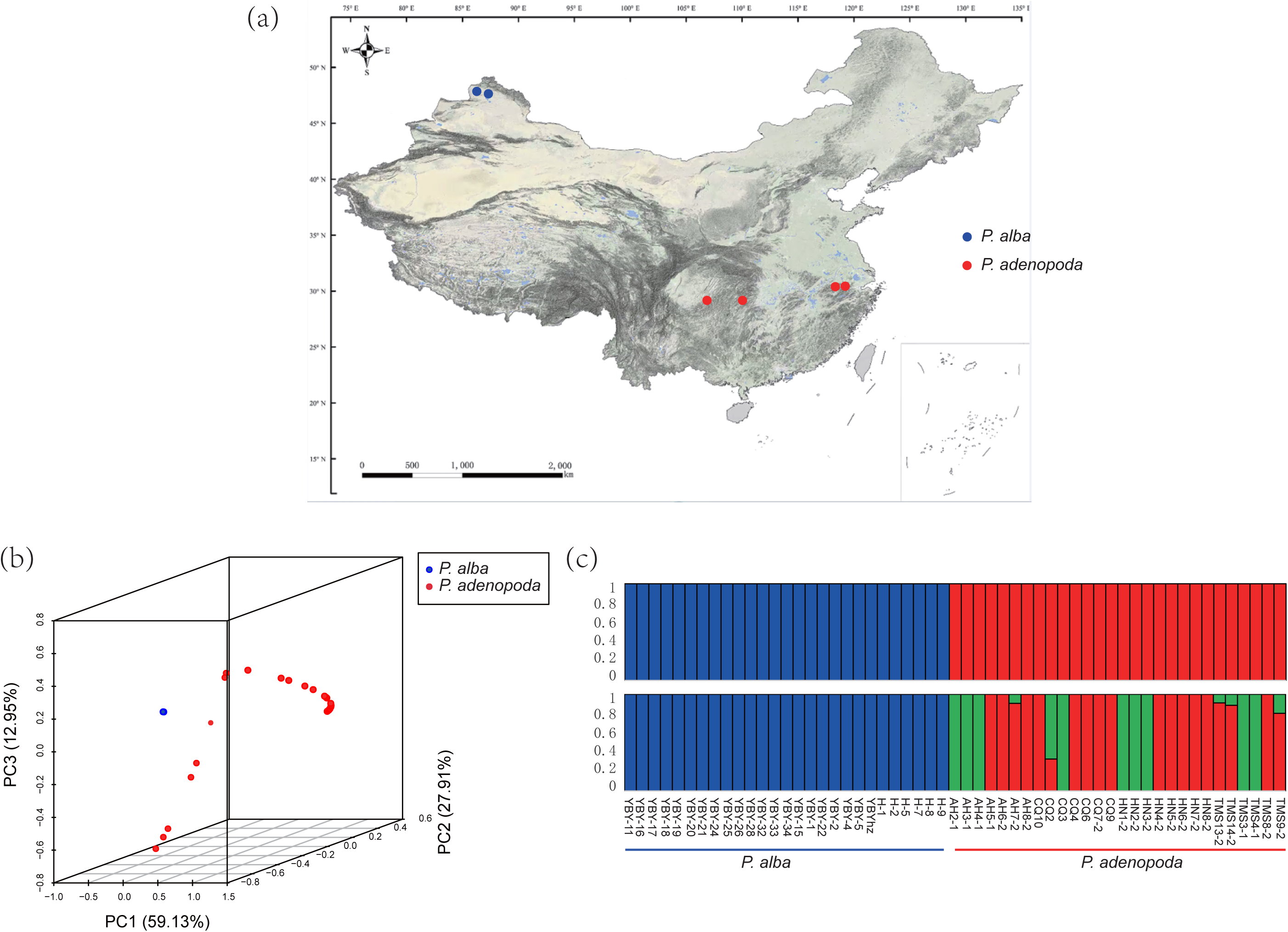
Sample collection and population structure analysis of 27 *P. alba* and 28 *P. adenopoda*. (a) 28 *P. adenopoda* individuals (red) were collected in Anhui, Chongqing, Hunan and Tianmushan of China and 27 *P. alba* individuals (blue) were collected from the Altay area, Xinjiang (a) (b) Population genetic structure in the samples based on an analysis using NGSadmix based on genotype likelihoods. The y-axis represents the number of clusters, and the x-axis shows the name of each individual. (c) Results from a principal component analysis (PCA) on the genetic covariance matrix for all individuals of *P. alba* (blue circles) and *P. adenopoda* (red circles).

We used *fastsimcoal2*.*6* (Excoffier et al., 2013), which is a continuous-time coalescence simulator assuming arbitrarily complex evolutionary scenarios (Excoffier and Foll, 2011), to infer the evolutionary history of *P. alba* and *P. adenopoda*. 25 different divergence models were assessed (Figure S1; Table S2). All models began with the subdivision of an ancestral population into two derived groups and thereafter differed in the following ways: (I) divergence time, (II) the occurrence and duration of gene flow following population divergence, and (III) effective population size before and after differentiation. The most appropriate model was an isolation-migration-isolation model, where the separation of the two species involved three phases (Figure 2a). Table 1 provides exact parameter estimates for this model, including the divergence time, levels of gene flow and effective population sizes together with their associated 95% confidence intervals (CIs). Under the best fitting model, *P. alba* and *P. adenopoda* diverged from an ancestral population approximately 9.54 million years ago (Mya) (bootstrap range [BR]: 5-10 Mya), and this differentiation initially occurred in allopatry as there was no evidence for gene flow during the early stages of differentiation. From 1.15 Mya to 0.34 Mya there was evidence for secondary contact and weak asymmetric gene flow between the two incipient species, after which the two species returned to an allopatric state without gene flow. The population size of the two isolated species showed stepwise changes, with *P. alba* experiencing a population decline during in the final stages. The migration rate of *P. alba* to *P. adenopoda* per generation was 1.28×10^−6^ (7.49×10^−7^-2.33×10^−6^), and that from *P. adenopoda* to *P. alba* was 9.69×10^−6^ (7.45×10^−6^-1.40×10^−5^). This result is not surprising, as the geographical distance between the two species is large and their distributions are very scattered. The estimates of contemporary effective population size (*Ne*) of *P. alba* (*N*_*e-P. alba*_) and *P. adenopoda* (*N*_*e-P. adenopoda*_) were 19,551 (BR:18624.5-23509.0) and 31,333 (BR:30851.5-34287.8), respectively, both of which were larger than the effective population size of the common ancestor which was 11,506 (BR:3170.7-80404.5).

**Table 1.** Demographic parameter estimates for the best-supported model in Figure 2a.

| Parameters | Point estimation | 95% CI<sup>a</sup> |  |
| --- | --- | --- | --- |
|  |  | Lower bound | Upper bound |
| $N_{E-ANC}$ | 11506 | 3170.675 | 80404.47 |
| $N_{E-POP0}$ | 19551 | 18624.45 | 23508.97 |
| $N_{E-POP1}$ | 31333 | 30851.53 | 34287.77 |
| **MIG01** | $1.28 \times 10^{-6}$ | $7.49 \times 10^{-7}$ | $2.33 \times 10^{-6}$ |
| **MIG10** | $9.59 \times 10^{-6}$ | $7.45 \times 10^{-6}$ | $1.40 \times 10^{-5}$ |
| **TDIV01** | 22786 | 21118.95 | 27847.42 |
| **TDIV02** | 76468 | 50890.35 | 98329.9 |
| **TDIV03** | 635956 | 336770.3 | 668025.3 |

<sup>a</sup>Parameter bootstrap estimation obtained by performing parameter estimation from 100 simulated data sets based on the total maximum composite likelihood estimates displayed in the point estimation column, per likelihood is estimated from 100,000 simulations.

**Figure 2.**
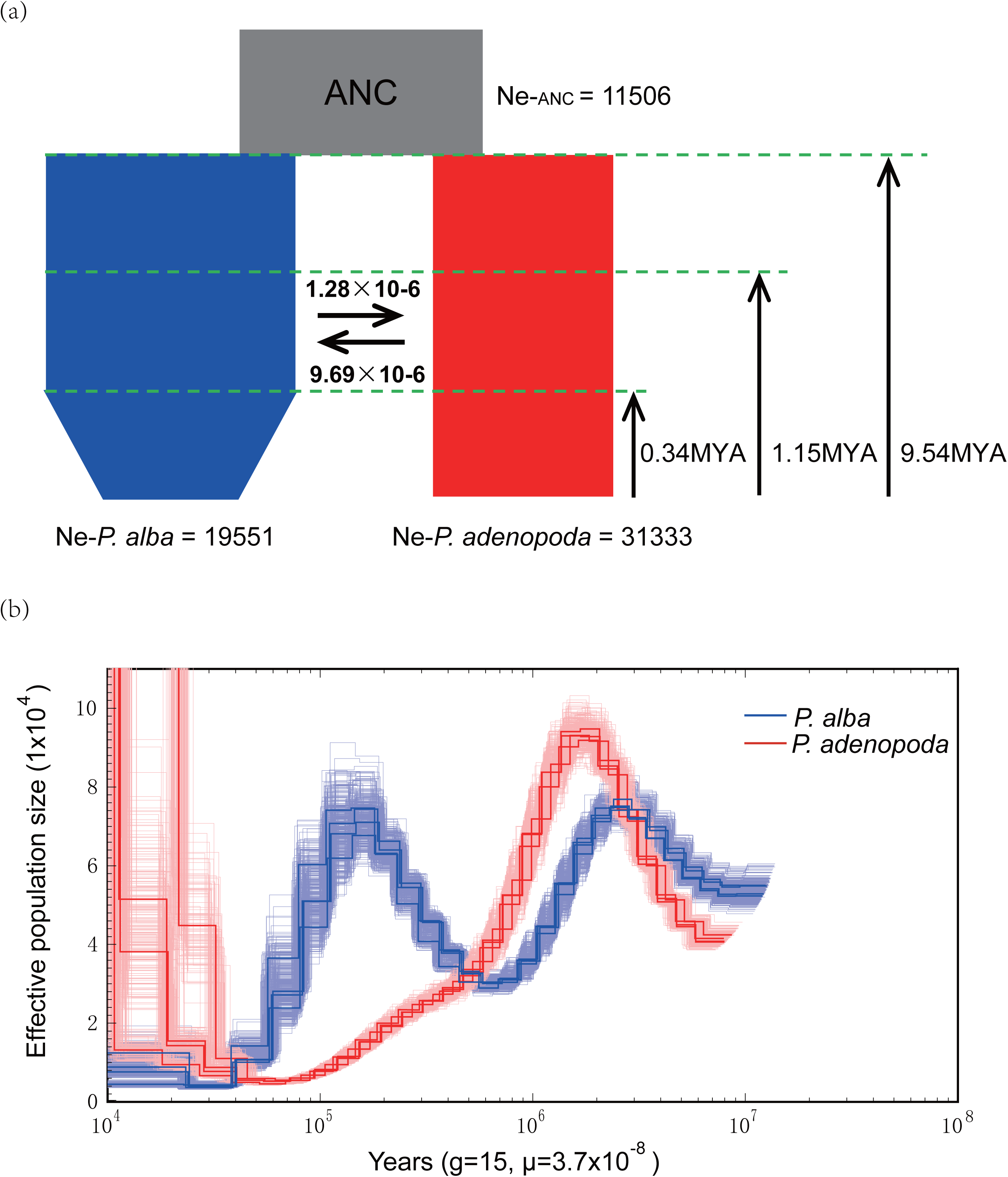
Demographic history of *P. alba* and *P. adenopoda*. (a) The most appropriate model inferred by *fastsimcoal2*.*6*. The ancestral population is colored gray while *P. alba* and the *P. adenopoda* are colored red and blue, respectively. The widths represents the relative effective population sizes (*N*e). Double-headed arrows represents the per generation gene flow between *P. alba* and *P. adenopoda*. All inferred demographic parameters are presented in Table 1. (b) Pairwise Sequentially Markovian coalescent (PSMC) estimates of the effective population size (*N*e) changes for *P. alba* (bold blue line) and *P. adenopoda* (bold red line) based on the inference from four phased haplotypes in each species, and performed 100 bootstrap replicates for four individuals (light-colored line respectively). Time scale on the x-axis is calculated assuming a neutral mutation rate per generation (µ) = 2.5×10-9 and generation time (g) = 15 years.

We also used the pairwise sequentially Markovian coalescent (PSMC) model (Li and Durbin, 2011) to reconstruct historical effective population size changes in the two species with a focus on diploid consensus sequences. Estimates of the effective population sizes of *P. alba* and *P. adenopoda* based on the PSMC model at the beginning of population divergence approximately 10 Mya were similar to those of their ancestral population obtained with *fastsimcoal2*.*6*. The two species experienced similar degrees of population expansion following divergence, possibly indicating successful adaptation to new niches (Figure 2b). The population size of *P. alba* then began to decline but rebounded with a second population expansion from 1.15-0.34 Mya. In contrast, the population of *P. adenopoda* has undergone considerable and continuous decline (Figure 2b).

### Genome-wide patterns of differentiation and identification of outlier regions

We used *F*_ST_, a relative measure of divergence (Charlesworth, 1998), to explore inter-specific genomic differentiation patterns in non-overlapping 10-kilobase (Kbp) windows (Figure 3). As shown in Figure 3, genetic differentiation varied widely across the genome with an average *F*_ST_ value of 0.542 and many windows showed substantially higher genetic differentiation between species. We identified 415 and 983 outlier windows which showed significantly (*P* < 0.05) high and low *F*_ST_ values, respectively (Figure 4a). By examining the genomic distribution of outlier windows, we determined that both highly and lowly differentiated regions were randomly distributed throughout the genome (Figure 3).

**Figure 3.**
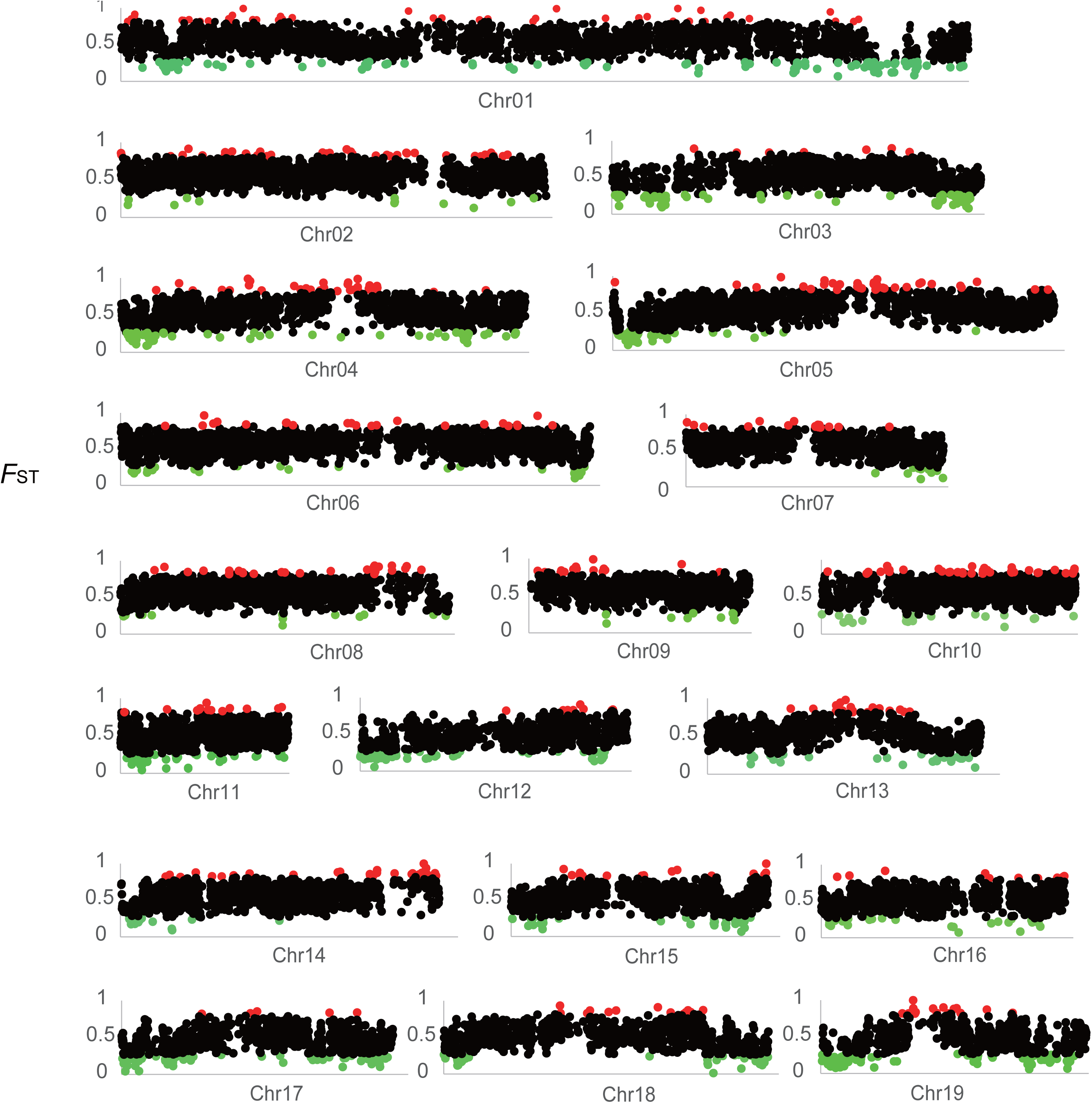
Genome-wide divergence. Chromosomal distribution of pairwise genetic divergence (*F*_ST_) between *P. alba* and *P. adenopoda* in 10-kb sliding windows. Windows with significantly high or low genetic differentiation are shown in red and green, respectively.

**Figure 4.**
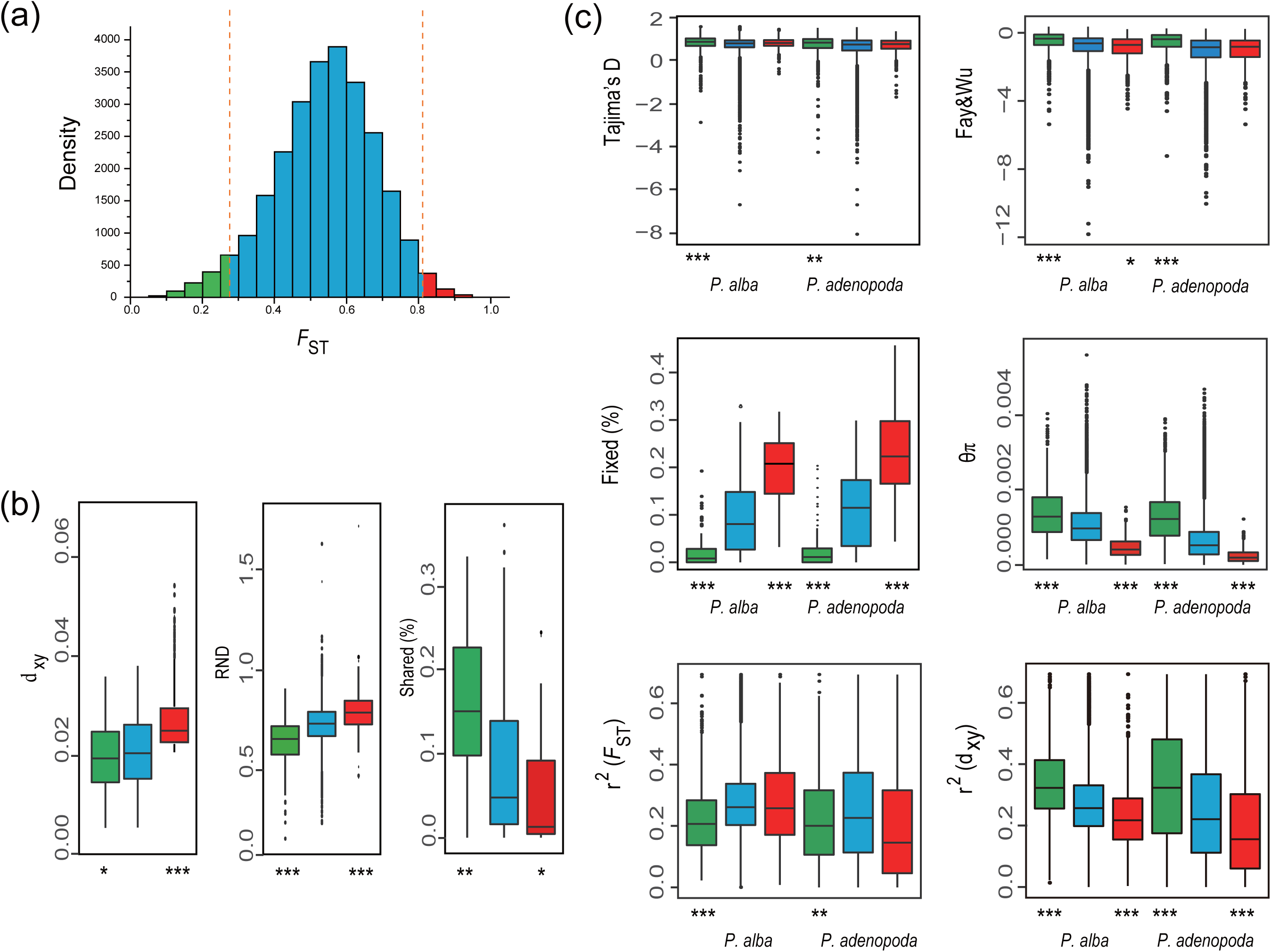
Identifying candidate outlier windows affected by factors of barriers to gene flow. (a) Distribution of genetic differentiation (*F*_ST_) between *P. alba* and *P. adenopoda*. The dotted line represents the threshold based on Z-scores to determine windows with high (red) (*P* > 0.05) or low (green) (*P* < −0.05) genetic differentiation. (b) d_xy_ (absolute measure of divergence), relative node depth (RND) and interspecific shared polymorphisms were compared between the genomic background and regions with significant high and low genetic differentiation. (c) Comparison of population genetic statistics between regions with significantly high (red) or low (green) genetic differentiation and the genomic background (blue) in *P. alba* and *P. adenopoda*. Statistics include Tajima’s D, Fay ʖWu’s H, the proportion of fixed differences caused by derived alleles, nucleotide diversity (θπ), and values of d_xy_ and *F*_ST_ calculated using the *r*^2^, respectively. Asterisks indicate significant differences between outliers and background genomic regions based on Mann-Whitney U tests (* *P*-value < 0.05; ** *P*-value < 1e-4; \*\*\**P*-value <2.2e-16).

Since *F*_ST_ is based on a comparison of intra- and inter-population diversity, any reduction in the former will increase *F*_ST_ (Nachman and Payseur, 2012). We used three other methods to quantify and compare inter-specific genetic differentiation between the two classes of outlier windows and the rest of the genome. The first was d_xy_, an absolute measure of divergence, which is the average number of pairwise differences between the two populations (Cruickshank and Hahn, 2014). The second was relative node depth (RND), which is d_xy_ divided by d_xy_ relative an outgroup species, in this case *P. trichocarpa*. This ratio corrects for possible variation in the mutation rate among genomic regions. The third was the proportion of ancestral polymorphisms shared by the two species. We found that d_xy_ and RND were significantly higher (Figure 4b; Table S3) and that the proportion of shared ancestral polymorphisms was significantly lower in the highly differentiated regions of the two species compared to the genomic background (Figure 4b; Table S3). These regions also contained a greater proportion of fixed differences, caused by the fixation of derived alleles in both species relative to the genome-wide average (Figure 4c; Table S3). Both species showed positive Tajima’s D values and negative Fay & Wu’s H values on average. The values of Tajima’s D and Fay & Wu’s H test showed no significant difference when comparing highly differentiated regions to the genomic background in the two species. Linkage disequilibrium (LD) in highly differentiated regions also showed no significant difference to the genomic background. However, when we divided the genome into three subsets based on d_xy_ values (high part, top 1000; medium and low part, bottom 1000), the d_xy_ values in the high and low parts showed significantly lower and higher value than the background in both the two species, respectively (Figure 4c; Table S3).

In contrast to the pattern found in highly differentiated regions, polymorphism in the lowly differentiated regions was higher. The value of Tajima’s D test and Fay & Wu’s H test were significantly higher for lowly differentiated regions than background regions. Compared with the genomic background, the proportion of polymorphisms shared between species was higher in lowly differentiated regions and the proportion of fixed differences was extremely low (Figure 4b, c; Table S3).

We also investigated the correlation between relative and absolute divergence between *P. alba* and *P. adenopoda* and we found a strong positive correlation between them (Spearman’s ρ= 0.623, *P*-value < 0.001).

A GO functional analysis of regions with high or low *F*_ST_ values was performed using the *P. trichocarpa* genome annotation. We assessed whether specific GO terms were significantly overexpressed using gene ontology (GO) assignments of these candidate genes. 293 and 599 genes were identified in the outlier windows showing high differentiation or low genetic differentiation, respectively (Table S4 and S5). We identified 41 GO categories showing significant enrichment, with the most prominent group (7 terms) associated with the term “reproduction “(*P* < 0.05) (Table S6). Genes related to reproduction and adaptation in highly differentiated regions between *P. alba* and *P. adenopoda* are listed on Tables S7 and S8.

## Discussion

### 1 Demography

We took advantage of genome-wide patterns of genetic diversity and differentiation to explore the evolutionary history of the two closely related species *P. alba* and *P. adenopoda* to study how the genomic heterogeneity in genetic diversity and differentiation is affected by various evolutionary forces.

Demographic modelling showed that *P. alba* and *P. adenopoda* diverged approximately 9.5 Mya (5 Mya-10 Mya), during the late Miocene. The climate of Asia has been strongly linked to the surface uplift of the Tibetan Plateau (Li et al., 2014a; Li et al., 2014b; Li et al., 2018; Zhang et al., 2000; Zhisheng et al., 2001). The Asian winter monsoon has appeared in some form since about 13.1 Mya (Fan et al., 2006). Rising vapour from the Indian Ocean cools, condenses and precipitates before it crosses the Tibetan Plateau, which results in drier air advancing to the northern plateau (Fang et al., 2019; Favre et al., 2015; Ge, 2006; Li et al., 2018). The combination of mountains and monsoons makes the climate in north-western China drier and the aridity reached its maximum approximately 9.6-8.0 Mya (Chen et al., 2019; Fan et al., 2007a; Mertz-Kraus et al., 2009; Zhisheng et al., 2001). From 5.3 Mya the climate has gradually became drier and/or cooler (Fan et al., 2007b). Furthermore, the uplift of the Pamir Plateau and the central plateau of Anatolia, coupled with the collision between the Pamir Plateau and the Tianshan Mountains, further blocked the transport of water vapour from the winter westerly belt to the east ∼5 Mya (Li et al., 2018; Meijers et al., 2018; Shen et al., 2018; Thompson et al., 2015). Approximately 2-3 Mya, the climate was still very dry and cold because north-western China received only the water vapour in the westerly belt at that time (Caves et al., 2015; Li et al., 2017; Zhuang et al., 2014). Approximately 1.80-1.10 Mya, the paleotemperature was approximately 7.5-13 °C (Yang and Ma, 2019), which was not sufficient to cause contact between *P. alba* and *P. adenopoda*. In our demographic modelling this is evident from the lack of gene flow that occurred during the early stages of divergence of the two species (from 9.54 Mya to 1.15 Mya).

Our modelling detects a small amount of asymmetric gene flow between the two species ∼ 1.15-0.34 Mya. This time period was close to the appearance of the Mindel-Riss Interglacial (Jing and Liu, 1999; Wright, 1926), which also included several cold and warm periods (Cheng, 1996; Lindner, 1981; Peiyuan, 1989). During this period, the paleotemperature increased significantly and the annual average temperature reached 28 °C (Cheng, 1996; Peiyuan, 1989). A form of subtropical forest or forest grassland environment was present during this time period and the climate was much warmer and wetter than in modern times. Based on the results of the PSMC simulation, the population of *P. alba* experienced exponential size changes at this stage. From ∼0.34 Mya to the present, gene flow between *P. alba* and *P. adenopoda* has been interrupted again and this period corresponds to the Riss Glacial (Jing and Liu, 1999; Kukla, 2005).

### 2 Heterogeneous differentiation across the genome

Relative divergence measures, such as *F*_ST_, are based on a comparison of within-population diversity to between-population diversity (Charlesworth, 1998; Cruickshank and Hahn, 2014). Absolute divergence is usually measured as d_xy_, the average number of pairwise differences between alleles sampled from two populations (Nachman Michael and Payseur Bret, 2012), and variation in d_xy_ (Ma et al., 2018) can be caused by variation in levels of ancestral polymorphism or substitution rates (Cruickshank and Hahn, 2014). A more comprehensive understanding of heterogeneity in genomic differentiation requires the use of both relative (influenced by within-population variation) and absolute (not influenced by within-population variation) measures of sequence divergence (Cruickshank and Hahn, 2014; Wolf and Ellegren, 2016). Any type of selection that reduces levels of linked neutral diversity will lead to increased estimates of relative divergence (such as *F*_ST_) (Charlesworth, 1998; Charlesworth et al., 1997; Cruickshank and Hahn, 2014). However, the increased *F*_ST_ seen in such regions does not always indicate a reduction in actual gene flow. In other words, there is a redistribution of alleles in these regions. Therefore, the value of d_xy_ in these regions does not necessarily increase. In contrast, if barriers to gene flow are the main driving force of heterogeneous differentiation across the genome, both absolute and relative divergence measures should be high because the redistribution of alleles in these regions is prevented while the remainder of the genome is homogenized by gene flow.

Most studies have implicated natural selection as the primary cause for genomic heterogeneous differentiation and even for speciation (Cruickshank and Hahn, 2014; Nosil and Feder, 2012). Selection affects population differentiation by reducing polymorphism levels at sites under selection (Cruickshank and Hahn, 2014). The size of differentiated regions is related to the intensity of selection and the stage of species divergence and the highly differentiated region may represent genes associated with reproductive isolation (Wu, 2001). However, barriers to gene flow, such as chromosomal rearrangements and genetic incompatibilities can also lead to heterogeneous differentiation of genomes (Faria and Navarro, 2010; Navarro et al., 1997; Navarro and Ruiz, 1997; Rieseberg, 2001). Barriers to gene flow inhibits allele exchange in such regions and in linked surrounding regions eventually leads to elevated genetic differentiation. If such highly differentiated regions contain genes associated with reproductive isolation, speciation may eventually occur (Cruickshank and Hahn, 2014; Wu, 2001). Our results found that absolute divergence (d_xy_) were strongly positively correlated with relative divergence (*F*_ST_) and this finding implies that barriers to gene flow have played a major role in generating the heterogeneous differentiation between *P. alba* and *P. adenopoda*, although natural selection cannot be thoroughly ruled out.

In this study, we Z-transformed the *F*_ST_ distributions to identify regions showing high, medium and low genetic differentiation across the genome. We evaluated multiple population genetic parameters to understand the mechanisms generating the heterogeneous genomic differentiation seen between the two species. Overall the values of Tajima’s D in the two species were positive and Fay & Wu’s H were negative, suggesting that recent population contractions have taken place (Fay and Wu, 2000; Holliday et al., 2010; Tajima, 1989), consistent with the PSMC results (Figure 2b). Both of the two neutrality tests (Tajima’s D and Fay & Wu’s H) showed no difference for the highly differentiated regions in either of the two species, suggesting that selection hasn’t been a the major factor in the maintenance of highly differentiated regions.

The extremely high correlation between relative and absolute divergence highlights the significant effects of barriers to gene flow in generating the heterogeneous differentiation landscape between the two species. Furthermore, when we divided the whole genome into three parts by d_xy_ values we found that the high part showed significantly lower correlation coefficients (*r*^2^) value than the genomic background, further supporting the conclusion that barriers to gene flow play an important role in shaping the genome-wide divergence of the two species (Ellegren et al., 2012; Geraldes et al., 2011). That may because when the foreign DNA fragments were introduced by introgression, the observed value of correlation coefficients (*r*^2^) increased in related regions, but not for the regions of barriers to gene flow because the foreign DNA fragment cannot be integrated into the genome. Compared to the genomic background, highly differentiated regions also contained an excess of derived fixed differences and a smaller proportion of shared polymorphisms. In addition, the nucleotide diversity (θπ) in highly differentiated regions was significantly greater than that of the genomic background, consistent with views outlined above. This also suggests that barriers to gene flow plays an important role in maintaining differentiation reproductive isolation in both species.

### 3 Reproductive isolation and speciation

Reproductive isolation is a consequence of prezygotic and postzygotic barriers as well as their potentially complex interactions (Rieseberg and Willis, 2007; Widmer et al., 2009). Prezygotic isolation, due to reduced probabilities for meeting (spatial or temporal isolation), mating (sexual isolation) or successful fertilization (homogametic sperm preferences or sperm-egg incompatibilities), has been found to be a greater contributor to the total isolation between species than postzygotic isolation (Qvarnström and Bailey, 2009).

Postzygotic isolation can be caused by intrinsic and extrinsic factors, leading to phenomena such as hybrid inviability and sterility and ecological and behavioural sterility, respectively (Widmer et al., 2009). *P. tomentosa*, one of the most widely distributed species in China, is a natural hybrid between *P. alba* and *P. adenopoda* (Li Kuan-yu, 1997; Wang et al., 2014) and while *P. tomentosa* has strong adaptability is has poor fertility and high abortion rates (Wang et al., 2019). A recent study using more than 200 individuals of *P. tomentosa* found no progeny resulting from backcrossing or selfing. Furthermore, some studies have found that *P. tomentosa* exhibits unequal numbers of univalents and a decreased degree of synapsis during meiosis (Kang, 2001). A larger percentage of univalents occurs at diakinesis and metaphase I and at anaphase I and telophase I lagging chromosomes are frequently observed (Kang et al., 1999). This finding is consistent with the conclusion of this study that heterogeneous genomic differentiation caused by barriers to gene flow is an important force in speciation. All of the above results confirmed that *P. alba* and *P. adenopoda* are subject to postzygotic isolation and imply that barriers to gene flow have played an important role in establishing and maintaining reproductive isolation between the two species.

There are several reasons for postzygotic barriers to gene flow, such as chromosomal rearrangements or gene incompatibilities (King and Templeton, 1993; Noor et al., 2001). Structural changes in the genome, including deletions, insertions, duplications, inversions and translocations, alter the genome organization of individuals and may also contribute to reproductive isolation (Wolf and Ellegren, 2016). Genomic incompatibilities might result from chromosomal rearrangements, which lead to mis-segregation during meiosis in hybrids, or from epistatic interactions that act as loss-of-function alleles in hybrid backgrounds (Lynch and Force, 2000). These factors are often accompanied by a decrease in hybrid inviability, sterility and/or even mortality (Barbash et al., 2003; Orr, 1989; Orr and Coyne, 1989; Schumer et al., 2014). To distinguish to what extent chromosomal rearrangements and gene incompatibilities contribute to postzygotic isolation, high-quality genomes for the two species must be assembled in the future.

To further categorise genes located in outlier regions, we performed gene ontology (GO) analysis in order to identify gene ontology terms that were enriched in these gene sets. We identified 41 categories that showing significant enrichment and the most prominent group (7 terms) was associated with the word “reproduction”. The seven genes belonging to this category include *CDC*2,*CUL*1,*emb*2746,*AFO,HK*2, *TFL*2 and *EDA*8, several of which are known to affect cell division, pollen rejection pathways, embryogenesis, flowers/spike into seedlings and reproductive habits, plant germ cell development during meiosis, transition from photoperiod regulation to reproductive growth and the early development of plant embryos and pollen tubes (Table S9). These findings support the hypothesis that highly differentiated regions contain an excess of reproduction-related genes. The gene ontology analysis also identified three genes involved in seed germination and a gene for drought resistance. *RCAR*3 and *GCR*1 regulated seed germination and early seedling development while *CPK*23 is involved in drought- and salt stress-induced calcium signaling cascades (Lim et al., 2014; Colucci et al., 2002; Ma et al., 2007). These genes may therefore be signatures of differential adaptation in the two species, *P. alba* and *P. adenopoda*.

## Conclusion

The uplift of the Tibetan Plateau and the Tianshan mountains led to drastic climate change in the northwest of China, which triggered the genetic differentiation of the two species. We observed that the absolute divergence (d_xy_) is significantly positive correlated with relative divergence (*F*_ST_), indicating that barriers to gene flow have played an important role in maintaining differentiation and reproductive isolation in these regions. By annotating genes in highly differentiated regions, we found that there are several genes associated with reproductive isolation and to adaptation to novel environments.

## Materials and Methods

### Population samples, sequencing and mapping

We sampled 27 individuals of *P. alba* and 28 individuals of *P. adenopoda* (Figure 1a and Table S1). We extracted genomic DNA from leaves of each individual using a DNA extraction kit (Aidlab, Beijing, China). Paired-end (PE) reads were constructed according to the Illumina library preparation protocol and sequenced on an Illumina HiSeq 2000 platform (Illumina, San Diego, CA). The sequencing coverage target for all samples was 25×. The data that support the findings of this study have been deposited into CNGB Sequence Archive (Guo et al., 2020) of CNGBdb with accession number CNP0001249 (http://db.cngb.org/cnsa/project/CNP0001249/reviewlink/). We obtained cleaned sequencing data from the sequencing company which we then mapped to the *P. trichocarpa* reference genome (v3.0) (Tuskan et al., 2006) using bwa-0.7.15 with default settings. We then performed PCR duplication removal on BAM files using MarkDuplicates from the Picard toolkit. Only reads with the highest summed base quality were used for downstream analyses.

### Filtering sites

We used several filtering steps to rule out errors that could be caused by paralogous or repeated DNA sequences before using the variant and genotype calls. First, we removed sites with a very low (<13 reads for each sample per species) or high (>100 reads for each sample per species) read coverage after examining the empirical distribution of reads. Second, since zero-mapped mass fractions were assigned to reads that could have been mapped to multiple genomic locations, we removed sites containing more than 20 such reads in all samples of each species. Third, we removed sites that overlapped with known repetitive elements identified using RepeatMasker (Chen, 2004; Tarailograovac and Chen, 2009). After these filtering steps, 48.6% of the sites in the genome were used for downstream analysis.

### SNP and genotype calling

Although ANGSD has been shown to be superior to the genotype calling methods in SAMtools and GATK (Depristo et al., 2011; Li et al., 2009), we used two complementary methods for SNP and genotype calling: i) Direct estimation of the genotype without calling in ANGSD v0.602 (Korneliussen et al., 2014). We considered only reads with a minimum mapping quality of 30 and bases with a minimum quality score of 20. For all the filtered sites in these two species, we defined the allele that was identical to that found in the *P. trichocarpa* reference genome as the ancestral allele. Based on the SAMtools genotype likelihood model for all sites, we used -doSaf to calculate the site allele frequency likelihood and then used -realSFS for maximum likelihood estimation of the expanded SFS using the expectation maximization (EM) algorithm (Su et al., 2011). Then, we calculated several population genetic statistics based on the global SFS, such as Fay & Wu’s H; the proportions of shared, private and fixed polymorphisms; and population structure parameters. ii) Multi-sample SNP and genotype calls were performed using HaplotypeCaller and GenotypeGVCFs in GATK by performing a series of filtering steps on the output VCF file. We also ran the a number of filtering steps to decrease false positives from SNP and genotype calls. We deleted SNPs in regions that did not pass all previous screening conditions and SNPs with more than two alleles in both species. Furthermore, if a genotype’s quality (GQ) was less than 10, it was designated as a missing genotype, and then the SNPs with more than two missing genotypes in each species were filtered out.

### Population structure

The NGSadmix program (Skotte et al., 2013) was used to infer population genetic structure by selecting only sites with less than 10% missing data. Based on the output from ANGSD, it was possible to infer individual admixture proportions and estimate the frequency of different ancestral populations based on genotype likelihoods. We utilized the SAMtools model (Li et al., 2009) in ANGSD to determine genotype likelihood and then generated a Beagle file of the genome subset using a likelihood ratio test (Su et al., 2011). The number of genetic clusters K was predetermined and varied from 2 to 4 and the maximum iteration of the EM algorithm was set to 10,000.

Similarly, the sample allele frequency likelihoods generated in ANGSD were used to perform a PCA to visualize inter-individual genetic relationships included possible sequencing errors and all uncertainties in the genotype calls (Fumagalli et al., 2014). The expected covariance matrix of each individual in the two species was calculated according to the genotypic posterior probability of all the filtered sites.

### Demographic history

We used *fastSimcoal2* (ver 2.6.0.3-14.10.17) (Excoffier et al., 2013) to infer the demographic history associated with the speciation of *P. alba* and *P. adenopoda* based simulations on a cluster. A two-dimensional joint SFS (2D-SFS) was constructed using the posterior probability from ngsTools for the frequency of the sample allele. We used 100,000 joint simulations to estimate the expected 2D-SFS and log probabilities for a set of demographic parameters in each model. The global maximum likelihood estimates for each model were obtained by 50 independent runs. We compared the models based on the maximum likelihood of 50 independent runs using the Akaike information criterion (AIC) and Akaike’s evidence weights. We chose the model with the largest Akaike’s weight value as the best model. We assumed that the mutation rate of poplar was 2.5×10^−9^ per year and that the generation time was 15 years when converting estimates to years and individual units. The CI of the best model was obtained from 100 parametric bootstrap samples, with 50 independent runs for each sample. We then used the PSMC method (Li and Durbin, 2011) to estimate historical changes in the population size of the two species. We used beagle.08Jun17.d8b.jar to phase and impute all of the sites within each species before analysis. The scaled time and population size were converted to actual time and size using a 15-year generation time and 2.5 × 10^−9^ mutations per nucleotide per year.

### Genome-wide patterns of differentiation

We divided the genome into 39,406 non-overlapping 10kbp windows for studying the pattern of genomic differentiation between the species. After all filtering steps detailed above, a window was used for downstream analyses if it included at least 1,000 bases. We also removed windows where the number of variable sites were less than 10. *F*_ST_ was calculated using the software VCFtools (v0.1.13) (Danecek et al., 2011) to estimate the degree of genetic differentiation between species at each site.

### Z-transformed *F*_ST_ values to find outlier regions

We used VCFtools (v0.1.13) (Danecek et al., 2011) to calculate the relative measure of divergence *F*_ST_ and then used the Origin 8 software to perform Z-transformation *F*_ST_. We chose to set the thresholds at Z(*F*_ST_)>1.96 and Z(*F*_ST_)<-1.96 as high and low-differentiated regions and 415 and 983 outlier windows were classified as highly and lowly differentiated regions, respectively. We compared two outlier windows (highly differentiated and lowly differentiated regions) with the rest of the genome by analysing additional population genetic statistics of the two species. First, we calculated θπ, Tajima’s D and Fay & Wu’s H according to the sample allele frequency likelihood of the non-overlapping 10-Kbp windows in ANGSD. Furthermore, we estimated and compared LD levels and recombination rates based on population size. We used VCFtools (Danecek et al., 2011) to calculate the correlation coefficients (*r*^2^) between SNPs with a distance greater than 1 Kbp to evaluate the level of LD in each 10-Kbp window. Finally, we used the ngsStat (Fumagalli et al., 2014) program to calculate several other genetic differentiation parameters: (1) the proportion of fixed differences caused by the fixation of derived alleles in *P. alba* and *P. adenopoda*, using *P. trichocarpa* as an outgroup; (2) the proportion of shared polymorphisms among species at all segregating sites; (3) the absolute measure of divergence (d_xy_) based on the posterior probability of the sample allele frequency for each locus and then averaged for each 10-Kbp window; and (4) RND, calculated by dividing the d_xy_ between *P. alba* and *P. adenopoda* by the d_xy_ between the two poplars and *P. trichocarpa*. For this process, we used one-sided Wilcoxon ranked-sum tests to detect significant differences between values in outlier windows and the genome-wide mean of all population genetic statistics.

### GO enrichment

To determine if any functional gene classes were overexpressed in the outlier regions with barriers to gene flow, we performed functional enrichment analysis of GO terms using Fisher’s exact test (https://www.omicshare.com/tools/Home/Soft/gogsea). This analysis excluded GO groups with fewer than two genes and further corrected the p-value from Fisher’s exact test using the Benjamini-Hochberg false discovery rate (Benjamini and Hochberg, 1995). GO terms with a modified *P*-value <0.05 were considered significantly enriched.

## Acknowledgments

We thank Dong Wang, Aiguo Duan, Wenhao Shao, Sirong Yi, Zhonghui Shi, Bin Zhang for some sample collection. We thank Prof. Jing Wang in Sichuan University, Associate Professor. Fumin Zhang, Dr. Zhe Cai and Chunyan Jing in the State Key Laboratory of Systematic and Evolutionary Botany, Institute of Botany, Chinese Academy of Sciences, Beijing, China, for their valuable suggestion on data analysis. This work was supported by China National GeneBank (CNGB).

