## Supplemental Table 1 for "Barriers to gene flow play an important role in miantaining reproductive isolation between two closely related *Populus* (Salicaceae) species"

**Table S1. Overview informatio of Illumina re-sequencing data per sample**

| Sample ID | Location | Latitude | Longitude | Mapping rate (%) | Mean Coverage |
|---|---|---|---|---|---|
| ***P.alba*** | | | | | |
| *P.alba*1 | Aletai | 47.5669 | 87.2533 | 95.23% | 23.432 |
| *P.alba*2 | Aletai | 47.3483 | 87.8697 | 95.29% | 23.953 |
| *P.alba*4 | Aletai | 47.8562 | 86.5721 | 95.21% | 20.471 |
| *P.alba*5 | Aletai | 47.8330 | 86.6641 | 95.08% | 23.772 |
| *P.alba*11 | Aletai | 47.6919 | 86.9401 | 94.02% | 25.479 |
| *P.alba*15 | Aletai | 47.6717 | 86.9699 | 94.60% | 21.449 |
| *P.alba*16 | Aletai | 47.6484 | 86.9746 | 94.54% | 27.622 |
| *P.alba*17 | Aletai | 47.6235 | 87.0252 | 93.87% | 24.601 |
| *P.alba*18 | Aletai | 47.6080 | 87.0711 | 94.63% | 25.185 |
| *P.alba*19 | Aletai | 47.5769 | 87.1332 | 94.49% | 23.165 |
| *P.alba*20 | Aletai | 47.5987 | 87.055 | 94.47% | 26.407 |
| *P.alba21* | Aletai | 47.4538 | 87.5196 | 94.61% | 20.246 |
| *P.alba*22 | Aletai | 47.4398 | 87.6599 | 94.18% | 26.869 |
| *P.alba*24 | Aletai | 47.4210 | 87.7288 | 94.50% | 24.901 |
| *P.alba*25 | Aletai | 47.3367 | 88.0025 | 94.09% | 23.821 |
| *P.alba*26 | Aletai | 47.4117 | 87.7174 | 93.96% | 21.650 |
| *P.alba*28 | Aletai | 47.5068 | 87.3218 | 94.09% | 18.222 |
| *P.alba*32 | Aletai | 47.3444 | 87.8892 | 94.43% | 20.958 |
| *P.alba33* | Aletai | 47.4911 | 87.3347 | 94.67% | 24.890 |
| *P.alba*34 | Aletai | 47.3914 | 87.7817 | 94.20% | 20.117 |
| *P.alba*H-1 | Aletai | 47.9011 | 86.1904 | 94.14% | 23.788 |
| *P.alba*H-2 | Aletai | 47.9073 | 86.0799 | 94.06% | 23.765 |
| *P.alba*H-5 | Aletai | 47.9274 | 86.0087 | 94.07% | 26.171 |
| *P.alba*H-7 | Aletai | 47.9938 | 85.6959 | 94.12% | 22.549 |
| *P.alba*H-8 | Aletai | 48.0185 | 85.6155 | 93.90% | 26.838 |
| *P.alba*H-9 | Aletai | 48.0324 | 85.6844 | 94.18% | 25.803 |
| YBYhz | Aletai | 48.0216 | 85.6178 | 94.71% | 28.777 |
| Mean | | | | 94.42% | 23.885 |
| ***P. adenopoda*** | | | | | |
| AH2-1 | Anhui | 30.2428 | 118.4423 | 95.17% | 22.644 |
| AH3-1 | Anhui | 30.2435 | 118.50 | 95.55% | 25.427 |
| AH4-1 | Anhui | 30.251 | 118.4645 | 95.62% | 33.695 |
| AH5-1 | Anhui | 30.2449 | 118.5827 | 95.43% | 27.740 |
| AH6-2 | Anhui | 30.2539 | 119.0318 | 94.90% | 23.745 |
| AH7-2 | Anhui | 30.2718 | 118.5722 | 94.96% | 24.329 |
| AH8-2 | Anhui | 30.2736 | 118.545 | 95.02% | 26.421 |
| CQ2 | Chongqing | 29.0812 | 107.0663 | 95.09% | 16.917 |

| | | | | | |
|---|---|---|---|---|---|
| CQ3 | Chongqing | 29.0787 | 107.0696 | 95.11% | 20.459 |
| CQ4 | Chongqing | 29.0771 | 107.0722 | 94.72% | 22.153 |
| CQ6 | Chongqing | 29.0755 | 107.0776 | 94.98% | 25.177 |
| CQ7-2 | Chongqing | 29.0716 | 107.0829 | 94.66% | 22.865 |
| CQ9 | Chongqing | 29.0691 | 107.0870 | 95.19% | 31.182 |
| CQ10 | Chongqing | 29.0677 | 107.0899 | 95.03% | 25.399 |
| HN1-2 | Hunan | 29.0664 | 110.2319 | 95.07% | 36.983 |
| HN2-2 | Hunan | 29.0678 | 110.2315 | 95.32% | 25.011 |
| HN3-2 | Hunan | 29.0697 | 110.2322 | 94.74% | 20.063 |
| HN4-2 | Hunan | 29.0675 | 110.2178 | 94.95% | 26.383 |
| HN5-2 | Hunan | 29.0711 | 110.2317 | 95.24% | 22.541 |
| HN6-2 | Hunan | 29.0711 | 110.2294 | 95.34% | 25.988 |
| HN7-2 | Hunan | 29.0664 | 110.2317 | 95.48% | 24.281 |
| HN8-2 | Hunan | 29.0642 | 110.2306 | 95.41% | 25.958 |
| TMS13-2 | Zhejiang | 30.3459 | 119.4460 | 94.77% | 24.012 |
| TMS14-2 | Zhejiang | 30.3456 | 119.4453 | 94.90% | 25.706 |
| TMS3-1 | Zhejiang | 30.3437 | 119.4439 | 95.78% | 19.825 |
| TMS4-1 | Zhejiang | 30.3450 | 119.4445 | 95.34% | 19.796 |
| TMS8-2 | Zhejiang | 30.3447 | 119.4430 | 95.05% | 22.956 |
| TMS9-2 | Zhejiang | 30.3444 | 119.4480 | 95.29% | 25.980 |
| Mean | | | | 95.15% | 24.773 |
