## Supplemental Table 2 for "Barriers to gene flow play an important role in miantaining reproductive isolation between two closely related *Populus* (Salicaceae) species"

### Table S2. Relative likelihood of the different models shown in Figure S1

| model | Max(log10(Lhoodi)[a] | No. Of parameters(d) | AICi[b] | Δi[b] | Model normalized relative likelihood(wi)b |
|---|---|---|---|---|---|
| Model1 | -85025483.6 | 4 | 391556830.1 | 681380.8686 | ~0 |
| Mode2 | -85103413.95 | 5 | 391915714.6 | 1040271.393 | ~0 |
| Model3 | -85056421.02 | 6 | 391699306.2 | 823858.9528 | ~0 |
| Model4 | -85033399.28 | 5 | 391593285.2 | 717843.9221 | ~0 |
| Model5 | -85019567.11 | 6 | 391529587.7 | 654144.4252 | ~0 |
| Model6 | -85048908.67 | 7 | 391664712.6 | 789263.3025 | ~0 |
| Model7 | -84982673.75 | 8 | 391359691.5 | 484250.2237 | ~0 |
| Model8 | -85023913.29 | 7 | 391549604.6 | 674153.3238 | ~0 |
| Model10 | -85043369.01 | 6 | 391639199.5 | 763756.2255 | ~0 |
| Model11 | -85022818.26 | 7 | 391544561.8 | 669112.5243 | ~0 |
| Model12 | -85025552.15 | 6 | 391557149.8 | 681706.553 | ~0 |
| Model13 | -85025440.78 | 7 | 391556638.9 | 681185.6752 | ~0 |
| Model14 | -85061928.28 | 8 | 391724672.1 | 849230.8223 | ~0 |
| Model15 | -84982165.92 | 9 | 391357354.8 | 481897.5801 | ~0 |
| Model16 | -85040874.24 | 8 | 391627714.6 | 752265.385 | ~0 |
| Model17 | -128561324.8 | 7 | 592046794 | 201171346.8 | ~0 |
| Model18 | -84929383.02 | 8 | 391114278.6 | 238827.3427 | ~0 |
| Model19 | -84974058.69 | 9 | 391320019.7 | 444566.4062 | ~0 |
| Model20 | -84967887.1 | 8 | 391291596.4 | 416147.1839 | ~0 |
| Model21 | -84962919.61 | 9 | 391268722.3 | 393273.0471 | ~0 |
| Model22 | -84877520.15 | 9 | 390875443.3 | 0 | 1 |
| Model23 | -84892761.84 | 12 | 390945639.8 | 70184.57637 | ~0 |
| Model24 | -84900737.75 | 12 | 390982370.3 | 106912.9993 | ~0 |
| Model25 | -84934178.07 | 12 | 391136368.6 | 260909.364 | ~0 |

[a] Based on the best likelihood among the 50 independent runs for each model (Figure S1).

[b] The calculation of $AIC_i$, $\Delta_i$ and $w_i$ are according to the methods shown in Excoffier *et al.* (2013).
