## Supplemental Table 3 for "Barriers to gene flow play an important role in miantaining reproductive isolation between two closely related *Populus* (Salicaceae) species"

**Table S3.** A comparison of the regions showing extreme genetic differentiation in *P. alba* and *P. adenopoda* with other genomic regions is presented (give the mean± standard deviation).

| **Parameters** | **Species** | **Regions displaying high differentiation** | **Regions displaying low differentiation** | **Background** |
|---|---|---|---|---|
| θπ | *P. alba* | 0.0005(±0.0003)*** | 0.0014(±0.0007)*** | 0.0011(±0.0006) |
| | *P. adenopoda* | 0.0002(±0.0002)*** | 0.0013(±0.0007)*** | 0.0007(±0.0005) |
| Tajima's D | *P. alba* | 1.2889(±0.5060) | 1.3644(±0.6763)*** | 1.1893(±0.6189) |
| | *P. adenopoda* | 1.1161(±0.6088) | 1.2098(±0.8052)** | 1.0416(±0.7827) |
| Fay & Wu's H | *P. alba* | -0.5939(±0.3420)* | -0.3731(±0.3552)*** | -0.5392(±0.3493) |
| | *P. adenopoda* | -0.7136(±0.4016) | -0.4640(±0.4417)*** | -0.7501(±0.4315) |
| $r^2(F_{ST})$ | *P. alba* | 0.3323(±0.2043) | 0.2568(±0.1531)*** | 0.3289(±0.1550) |
| | *P. adenopoda* | 0.2354(±0.2368) | 0.2634( ±0.1985)** | 0.3105(±0.2376) |
| $r^2$(Dxy) | *P. alba* | 0.2674(±0.1470)*** | 0.4213(±0.1776)*** | 0.3216(±0.1523) |
| | *P. adenopoda* | 0.2350( ±0.2104)*** | 0.4249(±0.2706)*** | 0.3048(±0.2341) |
| Fixed(%) | *P. alba* | 0.2146(±0.0887)*** | 0.0258(±0.0389)*** | 0.0999(±0.0851) |
| | *P. adenopoda* | 0.2535(±0.1122)*** | 0.0296(±0.0497)*** | 0.1211( ±0.0904) |
| Shared(%) | | 0.0517(±0.0680)* | 0.1695(±0.1009)** | 0.0908(±0.0965) |
| $F_{ST}$ | | 0.8594(±0.0396)*** | 0.2070(±0.0496)*** | 0.5496(±0.1194) |
| Dxy | | 0.0280(±0.0092)*** | 0.0212(±0.0083)* | 0.0220(±0.0085） |
| RND | | 1.2226(±0.2831)*** | 0.9213(±0.2040)*** | 1.0906(±0.1958） |

By Mann-Whitney U test, there were significant differences between the genome background and regions showing extreme genetic differentiation (* *P*-value < 0.05; ** *P*-value < 1e-4; ****P*-value <2.2e-16).
