## Supplemental Table 4 for "Barriers to gene flow play an important role in miantaining reproductive isolation between two closely related *Populus* (Salicaceae) species"

**Table S4**. List of genes located in a region of significantly high genetic differentiation between *P. alba* and *P. adenopoda*.

| Poplar gene | Best Arabidopsi hit | Synonyms | Annotated description |
|---|---|---|---|
| Potri.001G009600 | AT1G69770 | CMT3 | chromomethylase 3 |
| Potri.001G011500 | | | |
| Potri.001G011900 | AT3G07160 | GSL10 | glucan synthase-like 10 |
| Potri.001G012200 | AT5G13000 | GSL12 | glucan synthase-like 12 |
| Potri.001G048800 | AT1G15520 | ABCG40 | pleiotropic drug resistance 12 |
| Potri.001G049400 | AT2G32700 | LUH | LEUNIG_homolog |
| Potri.001G053200 | | | |
| Potri.001G065800 | AT5G13260 | | |
| Potri.001G083400 | AT4G24140 | | alpha/beta-Hydrolases superfamily protein |
| Potri.001G083900 | AT2G46660 | CYP78A6 | cytochrome P450, family 78, subfamily A, polypeptide 6 |
| Potri.001G092500 | AT5G53160 | RCAR3 | regulatory components of ABA receptor 3 |
| Potri.001G099400 | AT4G11280 | ACS6 | 1-aminocyclopropane-1-carboxylic acid (acc) synthase 6 |
| Potri.001G101300 | AT5G41360 | XPB2 | homolog of Xeroderma pigmentosum complementation group B 2 |
| Potri.001G117000 | AT1G12330 | | (1 of 1) PTHR31029:SF4 - F5O11.6 |
| Potri.001G118500 | AT4G12310 | CYP706A5 | cytochrome P450, family 706, subfamily A, polypeptide 5 |
| Potri.001G119000 | AT1G62790 | | Bifunctional inhibitor/lipid-transfer protein/seed storage 2S albumin superfamily protein |
| Potri.001G125900 | | | |
| Potri.001G190400 | | | |
| Potri.001G196500 | AT1G17640 | | RNA-binding (RRM/RBD/RNP motifs) family protein |
| Potri.001G198000 | AT1G54220 | | Dihydrolipoamide acetyltransferase, long form protein |
| Potri.001G211900 | | | |
| Potri.001G215700 | AT1G08420 | BSL2 | BRI1 suppressor 1 (BSU1)-like 2 |
| Potri.001G233600 | AT5G59990 | | CCT motif family protein |
| Potri.001G249900 | AT2G29660 | | zinc finger (C2H2 type) family protein |
| Potri.001G305400 | | | |
| Potri.001G305800 | | | |
| Potri.001G309650 | | | |
| Potri.001G324800 | | | |

| | | | |
|---|---|---|---|
| Potri.001G335300 | AT5G40780 | LHT1 | lysine histidine transporter 1 |
| Potri.001G340800 | | | |
| Potri.001G342700 | AT5G40550 | SGF29b | SGF29 tudor-like domain |
| Potri.001G352900 | | | |
| Potri.001G355200 | | | |
| Potri.001G361900 | AT4G37190 | | (1 of 1) PTHR13391 - TUBULIN-RELATED PROTEIN |
| Potri.001G400900 | AT1G53000 | CKS | Nucleotide-diphospho-sugar transferases superfamily protein |
| Potri.001G414700 | AT4G27310 | BBX28 | B-box type zinc finger family protein |
| Potri.002G000200 | | | |
| Potri.002G046300 | | | |
| Potri.002G055000 | | | |
| Potri.002G059200 | | | |
| Potri.002G070300 | AT5G42140 | | Regulator of chromosome condensation (RCC1) family with FYVE zinc finger domain |
| Potri.002G087300 | AT1G38065 | | O-fucosyltransferase family protein |
| Potri.002G087600 | AT1G77720 | PPK1 | putative protein kinase 1 |
| Potri.002G092500 | AT1G22150 | SULTR1;3 | sulfate transporter 1;3 |
| Potri.002G102700 | AT4G24390 | AFB4 | RNI-like superfamily protein |
| Potri.002G110900 | AT4G24470 | ZIM | GATA-type zinc finger protein with TIFY domain |
| Potri.002G111200 | AT4G24480 | | Protein kinase superfamily protein |
| Potri.002G115000 | AT1G58470 | RBP1 | RNA-binding protein 1 |
| Potri.002G118950 | | | |
| Potri.002G159300 | AT2G45880 | BAM7 | beta-amylase 7 |
| Potri.002G161700 | AT4G00310 | EDA8 | Putative membrane lipoprotein |
| Potri.002G164100 | AT5G12370 | SEC10 | exocyst complex component sec10 |
| Potri.002G176900 | | | |
| Potri.002G191000 | AT2G47160 | BOR1 | HCO3- transporter family |
| Potri.002G197200 | AT1G02640 | BXL2 | beta-xylosidase 2 |
| Potri.002G198500 | AT1G66530 | | Arginyl-tRNA synthetase, class Ic |
| Potri.002G202200 | AT1G02790 | PGA4 | polygalacturonase 4 |
| Potri.002G206100 | AT2G47770 | TSPO | TSPO(outer membrane tryptophan-rich sensory protein)-related |
| Potri.002G213300 | AT4G03080 | BSL1 | BRI1 suppressor 1 (BSU1)-like 1 |
| Potri.002G223100 | | | |
| Potri.002G224900 | AT2G32970 | RTL1 | |
| Potri.002G226700 | AT4G15417 | ACO3 | RNAse II-like 1 |
| Potri.002G229200 | AT2G05710 | GLR3.4 | aconitase 3 |
| Potri.002G230000 | AT1G05200 | | glutamate receptor 3.4 |

| | | | |
|---|---|---|---|
| Potri.002G233500 | AT1G10840 | TIF3H1 | translation initiation factor 3 subunit H1 |
| Potri.002G235700 | AT5G42390 | SPP | Insulinase (Peptidase family M16) family protein |
| Potri.003G050200 | AT1G79770 | EER4 | Protein of unknown function (DUF1677) |
| Potri.003G064100 | AT1G17440 | | Transcription initiation factor TFIID subunit A |
| Potri.003G130500 | AT1G63820 | | CCT motif family protein |
| Potri.003G151300 | AT5G51180 | | alpha/beta-Hydrolases superfamily protein |
| Potri.004G029200 | | | |
| Potri.004G046201 | | | |
| Potri.004G054900 | AT4G18570 | | Tetratricopeptide repeat (TPR)-like superfamily protein |
| Potri.004G079000 | AT5G52530 | | dentin sialophosphoprotein-related |
| Potri.004G090600 | | | |
| Potri.004G091200 | AT1G07790 | HTB1 | Histone superfamily protein |
| Potri.004G097900 | AT1G67030 | ZFP6 | zinc finger protein 6 |
| Potri.004G116500 | AT3G29770 | MES11 | methyl esterase 11 |
| Potri.004G121200 | AT5G15400 | | U-box domain-containing protein |
| Potri.004G123900 | AT3G28720 | | (1 of 1) PTHR31515:SF3 - GENOMIC DNA, CHROMOSOME 3,BAC CLONE: T19N8 |
| Potri.004G126100 | AT5G39410 | | Saccharopine dehydrogenase |
| Potri.004G127400 | AT5G39570 | | (1 of 2) PTHR33971:SF1 - GENOMIC DNA, CHROMOSOME 3,P1 CLONE:MRI12 |
| Potri.004G128400 | AT3G28857 | PRE5 | basic helix-loop-helix (bHLH) DNA-binding family protein |
| Potri.004G129000 | | | |
| Potri.004G129200 | AT5G39830 | DEG8 | Trypsin family protein with PDZ domain |
| Potri.004G131800 | AT2G36160 | | Ribosomal protein S11 family protein |
| Potri.004G131900 | AT1G29990 | PFD6 | prefoldin 6 |
| Potri.004G133500 | AT3G48750 | CDC2 | cell division control 2 |
| Potri.004G133600 | AT5G08430 | | SWIB/MDM2 domain;Plus-3;GYF |
| Potri.004G133700 | AT5G08440 | | |
| Potri.005G109500 | AT4G35900 | FD | Basic-leucine zipper (bZIP) transcription factor family protein |
| Potri.005G126000 | AT4G36730 | GBF1 | G-box binding factor 1 |
| Potri.005G136600 | | | |
| Potri.005G139400 | AT4G36945 | | PLC-like phosphodiesterases superfamily protein |
| Potri.005G144700 | AT2G23140 | PUB4 | RING/U-box superfamily protein with ARM repeat domain |
| Potri.005G144800 | AT4G37440 | | |
| Potri.005G146800 | AT5G67500 | VDAC2 | voltage dependent anion channel 2 |

| | | | |
|---|---|---|---|
| Potri.005G148400 | AT4G37750 | ANT | Integrase-type DNA-binding superfamily protein |
| Potri.005G152500 | | | |
| Potri.005G152800 | AT4G24470 | | GATA-type zinc finger protein with TIFY domain |
| Potri.005G153200 | AT1G09700 | HYL1 | dsRNA-binding domain-like superfamily protein |
| Potri.005G153800 | AT4G24520 | ATR1 | P450 reductase 1 |
| Potri.005G155300 | | | |
| Potri.005G156100 | AT2G01710 | | Chaperone DnaJ-domain superfamily protein |
| Potri.005G156500 | | | |
| Potri.005G158300 | AT5G10940 | ASG2 | transducin family protein / WD-40 repeat family protein |
| Potri.005G161700 | AT1G35210 | | |
| Potri.005G168800 | AT1G36050 | | Endoplasmic reticulum vesicle transporter protein |
| Potri.005G186500 | AT1G21450 | SCL1 | SCARECROW-like 1 |
| Potri.005G187300 | AT1G21460 | SWEET1 | Nodulin MtN3 family protein |
| Potri.005G254200 | | | |
| Potri.006G038000 | | | |
| Potri.006G065900 | | | |
| Potri.006G076300 | AT5G57230 | | Thioredoxin superfamily protein |
| Potri.006G123000 | AT5G03430 | | phosphoadenosine phosphosulfate (PAPS) reductase family protein |
| Potri.006G126700 | AT3G09810 | IDH-VI | isocitrate dehydrogenase VI |
| Potri.006G152500 | AT5G56970 | CKX3 | cytokinin oxidase 3 |
| Potri.006G153700 | AT3G57340 | | Heat shock protein DnaJ, N-terminal with domain of unknown function (DUF1977) |
| Potri.006G155400 | AT5G55740 | CRR21 | Tetratricopeptide repeat (TPR)-like superfamily protein |
| Potri.006G160800 | AT1G67290 | GLOX1 | glyoxal oxidase-related protein |
| Potri.006G179100 | AT1G73250 | GER1 | GDP-4-keto-6-deoxymannose-3,5-epimerase -4-reductase 1 |
| Potri.006G201100 | | | |
| Potri.006G213700 | AT3G53540 | TRM19 | (1 of 2) PTHR21726:SF49 - PHOSPHAT-IDYLINOSITOL N-ACETYGLUCOSA MINLYTRANSFERASE SUBUNIT P-LIKE PROTEIN |
| Potri.006G218000 | AT2G26250 | KCS10 | 3-ketoacyl-CoA synthase 10 |
| Potri.006G223600 | AT1G72820 | | Mitochondrial substrate carrier family protein |

| | | | |
|---|---|---|---|
| Potri.006G223900 | AT5G12030 | HSP17.6A | heat shock protein 17.6A |
| Potri.006G250600 | AT4G32440 | | Plant Tudor-like RNA-binding protein |
| Potri.007G000700 | AT3G20010 | | SNF2 domain-containing protein / helicase domain-containing protein / zinc finger protein-related |
| Potri.007G006900 | AT4G37730 | bZIP7 | basic leucine-zipper 7 |
| Potri.007G014400 | AT4G36160 | NAC076 | NAC domain containing protein 76 |
| Potri.007G043800 | AT4G36990 | HSF4 | heat shock factor 4 |
| Potri.007G060800 | AT5G66060 | | 2-oxoglutarate (2OG) and Fe(II)-dependent oxygenase superfamily protein |
| Potri.007G063200 | AT1G21630 | | Calcium-binding EF hand family protein |
| Potri.007G063600 | AT4G35890 | LARP1c | winged-helix DNA-binding transcription factor family protein |
| Potri.007G063700 | AT4G35870 | | early-responsive to dehydration stress protein (ERD4) |
| Potri.007G064000 | | | |
| Potri.007G064300 | AT2G17480 | MLO8 | Seven transmembrane MLO family protein |
| Potri.007G066900 | AT1G16520 | | |
| Potri.007G070501 | | | |
| Potri.008G045200 | AT5G13610 | | Protein of unknown function (DUF155) |
| Potri.008G101500 | AT3G06500 | A/N-InvC | Plant neutral invertase family protein |
| Potri.008G113400 | AT2G04038 | bZIP48 | basic leucine-zipper 48 |
| Potri.008G121300 | AT1G25420 | | Regulator of Vps4 activity in the MVB pathway protein |
| Potri.008G142600 | | | |
| Potri.008G144700 | AT2G02220 | PSKR1 | phytosulfokin receptor 1 |
| Potri.008G156800 | AT4G14746 | | |
| Potri.008G182700 | AT1G13180 | DIS1 | Actin-like ATPase superfamily protein |
| Potri.008G203400 | AT5G19140 | AILP1 | Aluminium induced protein with YGL and LRDR motifs |
| Potri.008G206000 | AT1G48270 | GCR1 | G-protein-coupled receptor 1 |
| Potri.008G206100 | | | |
| Potri.008G207500 | AT4G15790 | | |
| Potri.008G210200 | AT2G47820 | | (1 of 4) PTHR13859 - ATROPHIN-RELATED |
| Potri.008G213500 | AT3G16857 | RR1 | response regulator 1 |
| Potri.008G217300 | AT4G02570 | CUL1 | cullin 1 |
| Potri.009G004500 | AT1G07960 | PDIL5-1 | PDI-like 5-1 |
| Potri.009G006200 | AT2G28130 | | |
| Potri.009G010350 | AT5G60710 | | Zinc finger (C3HC4-type RING finger) family protein |

| | | | |
|---|---|---|---|
| Potri.009G010600 | AT4G27490 | RRP41L | 3'-5'-exoribonuclease family protein |
| Potri.009G015100 | AT5G60220 | TET4 | tetraspanin4 |
| Potri.009G016600 | AT5G22090 | | Protein of unknown function (DUF3049) |
| Potri.009G026800 | AT1G07720 | KCS3 | 3-ketoacyl-CoA synthase 3 |
| Potri.009G026900 | AT5G59730 | EXO70H7 | exocyst subunit exo70 family protein H7 |
| Potri.009G034900 | AT2G29140 | PUM3 | pumilio 3 |
| Potri.009G140200 | AT1G16210 | | (1 of 1) PTHR21680:SF0 - COILED-COIL DOMAIN-CONTAINING PROTEIN 124 |
| Potri.010G006300 | AT3G16785 | PLDP1 | phospholipase D P1 |
| Potri.010G028900 | AT4G15470 | | Bax inhibitor-1 family protein |
| Potri.010G029500 | AT5G19000 | BPM1 | BTB-POZ and MATH domain 1 |
| Potri.010G032900 | AT2G42770 | | Peroxisomal membrane 22 kDa (Mpv17/ PMP22) family protein |
| Potri.010G033900 | AT3G17900 | | |
| Potri.010G034300 | AT1G69850 | NRT1:2 | nitrate transporter 1:2 |
| Potri.010G034800 | | | |
| Potri.010G042300 | AT1G10657 | | Plant protein 1589 of unknown function |
| Potri.010G079300 | AT1G04220 | KCS2 | 3-ketoacyl-CoA synthase 2 |
| Potri.010G089700 | AT3G22380 | TIC | time for coffee |
| Potri.010G090900 | AT5G19330 | ARIA | ARM repeat protein interacting with ABF2 |
| Potri.010G099300 | AT1G75010 | ARC3 | GTP binding |
| Potri.010G105200 | | | |
| Potri.010G116000 | AT5G17350 | | (1 of 28) PF14009 - Domain of unknown function (DUF4228) (DUF4228) |
| Potri.010G128200 | AT1G25510 | | Eukaryotic aspartyl protease family protein |
| Potri.010G128600 | AT1G25540 | PFT1 | phytochrome and flowering time regulatory protein (PFT1) |
| Potri.010G129800 | | | |
| Potri.010G132700 | | | |
| Potri.010G136200 | AT1G68930 | | pentatricopeptide (PPR) repeat-containing protein |
| Potri.010G164500 | AT1G14000 | VIK | VH1-interacting kinase |
| Potri.010G165700 | | | |
| Potri.010G165800 | AT2G03260 | | EXS (ERD1/XPR1/SYG1) family protein |
| Potri.010G169200 | AT5G62460 | | RING/FYVE/PHD zinc finger superfamily protein |
| Potri.010G188700 | AT5G05660 | NFXL2 | sequence-specific DNA binding transcription factors;zinc ion binding |
| Potri.010G200200 | AT1G74260 | PUR4 | purine biosynthesis 4 |
| Potri.010G201400 | AT2G39710 | | Eukaryotic aspartyl protease family protein |
| Potri.010G223300 | AT3G54810 | BME3 | Plant-specific GATA-type zinc finger transc- |

| | | | |
|---|---|---|---|
| | | | ription factor family protein |
| Potri.010G238400 | | | |
| Potri.010G250800 | AT3G21180 | ACA9 | autoinhibited Ca(2+)-ATPase 9 |
| Potri.010G252000 | AT1G78880 | | Ubiquitin-specific protease family C19-related protein |
| Potri.010G254300 | | | |
| Potri.010G254500 | AT3G19980 | FYPP3 | flower-specific, phytochrome-associated protein phosphatase 3 |
| Potri.011G003400 | AT4G04740 | CPK23 | calcium-dependent protein kinase 23 |
| Potri.011G082500 | AT4G18375 | | RNA-binding KH domain-containing protein |
| Potri.011G085100 | AT1G30070 | | SGS domain-containing protein |
| Potri.011G086800 | AT3G20800 | | Cell differentiation, Rcd1-like protein |
| Potri.011G091900 | AT1G30330 | ARF6 | auxin response factor 6 |
| Potri.011G095600 | AT2G34555 | ATGA2OX3 | gibberellin 2-oxidase 3 |
| Potri.011G107800 | AT3G14860 | | NHL domain-containing protein |
| Potri.011G114200 | | | |
| Potri.011G129000 | | | |
| Potri.011G158100 | AT5G44410 | | FAD-binding Berberine family protein |
| Potri.011G163800 | | | |
| Potri.012G066000 | AT1G74240 | | Mitochondrial substrate carrier family protein |
| Potri.012G095800 | AT5G63420 | emb2746 | RNA-metabolising metallo-beta-lactamase family protein |
| Potri.012G100400 | AT5G63510 | GAMMA CAL1 | gamma carbonic anhydrase like 1 |
| Potri.012G105900 | AT5G50920 | CLPC1 | CLPC homologue 1Integrase-type DNA-binding superfamily protein |
| Potri.012G108500 | AT5G07310 | | |
| Potri.012G132400 | AT4G25420 | GA20OX1 | 2-oxoglutarate (2OG) and Fe(II)-dependent oxygenase superfamily protein |
| Potri.013G064600 | AT3G03750 | | SET domain protein 20 |
| Potri.013G074100 | AT5G17560 | | BolA-like family protein |
| Potri.013G083300 | AT3G10960 | AZG1 | AZA-guanine resistant1 |
| Potri.013G085600 | AT1G65220 | | ARM repeat superfamily protein |
| Potri.013G086600 | AT5G16490 | RIC4 | ROP-interactive CRIB motif-containing |
| Potri.013G087950 | AT1G65080 | | protein 4 Membrane insertion protein,OxaA/ YidC with tetratricopeptide repeat domain |
| Potri.013G088100 | AT1G31800 | CYP97A3 | cytochrome P450, family 97, subfamily A, polypeptide 3 |
| Potri.013G090400 | AT2G40740 | WRKY55 | WRKY DNA-binding protein 55 |
| Potri.013G092400 | AT4G10350 | NAC070 | NAC domain containing protein 70 |
| Potri.013G094950 | | | |
| Potri.013G101601 | AT1G22840 | CYTC-1 | CYTOCHROME C-1 |

| | | | |
|---|---|---|---|
| Potri.013G103551 | | | |
| Potri.014G032600 | | | |
| Potri.014G035500 | | | |
| Potri.014G044700 | | | |
| Potri.014G057700 | AT1G02065 | SPL8 | squamosa promoter binding protein-like 8 |
| Potri.014G066700 | AT2G45190 | AFO | Plant-specific transcription factor YABBY family protein |
| Potri.014G074200 | AT2G45660 | AGL20 | AGAMOUS-like 20 |
| Potri.014G084900 | AT2G45910 | | U-box domain-containing protein kinase family protein |
| Potri.014G097900 | | | |
| Potri.014G134601 | AT2G47890 | | B-box type zinc finger protein with CCT domain |
| Potri.014G163100 | AT2G04750 | | Actin binding Calponin homology (CH) domain-containing protein |
| Potri.014G164700 | AT5G35750 | HK2 | histidine kinase 2 |
| Potri.014G179500 | AT3G60150 | | Protein of unknown function (DUF498/ DUF598) |
| Potri.014G179600 | AT2G44530 | | Phosphoribosyltransferase family protein |
| Potri.014G180200 | AT1G08840 | emb2411 | DNA replication helicase, putative |
| Potri.014G180700 | AT1G45150 | | (1 of 1) PF13320 - Domain of unknown function (DUF4091) |
| Potri.014G180800 | AT5G42970 | COP8 | Proteasome component (PCI) domain protein |
| Potri.014G185300 | AT5G42710 | TRM30 | |
| Potri.014G189900 | AT3G24880 | | Helicase/SANT-associated, DNA binding protein |
| Potri.014G192200 | AT5G44010 | | (1 of 1) PF11107 - Fanconi anemia group F protein (FANCF) |
| Potri.014G192300 | AT5G44000 | | Glutathione S-transferase family protein |
| Potri.014G192500 | AT1G04050 | SUVR1 | homolog of SU(var)3-9 1 |
| Potri.014G192800 | AT1G04080 | PRP39 | Tetratricopeptide repeat (TPR)-like superfamily protein |
| Potri.014G196400 | AT5G42770 | | Maf-like protein |
| Potri.014G196900 | AT1G47330 | | CBS domain-containing protein with a domain of unknown function (DUF21) |
| Potri.015G039500 | AT1G73630 | | EF hand calcium-binding protein family |
| Potri.015G043100 | AT3G18210 | | 2-oxoglutarate (2OG) and Fe(II)-dependent oxygenase superfamily protein |
| Potri.015G043200 | AT1G17980 | PAPS1 | poly(A) polymerase 1 |
| Potri.015G056800 | AT5G05810 | ATL43 | RING/U-box superfamily protein |

| | | | |
|---|---|---|---|
| Potri.015G057700 | AT2G33040 | ATP3 | gamma subunit of Mt ATP synthase |
| Potri.015G071100 | | | |
| Potri.015G074200 | AT1G10560 | PUB18 | plant U-box 18 |
| Potri.015G100600 | | | |
| Potri.015G134600 | AT4G25420 | | 2-oxoglutarate (2OG) and Fe(II)-dependent oxygenase superfamily protein |
| Potri.015G145300 | AT4G20850 | TPP2 | tripeptidyl peptidase ii |
| Potri.015G147200 | AT4G09980 | EMB1691 | Methyltransferase MT-A70 family protein |
| Potri.016G017300 | AT3G21800 | UGT71B8 | UDP-glucosyl transferase 71B8 |
| Potri.016G029500 | AT3G56550 | | Pentatricopeptide repeat (PPR) superfamily protein |
| Potri.016G056800 | AT2G41660 | MIZ1 | Protein of unknown function, DUF617 |
| Potri.016G099700 | AT3G09310 | | (1 of 1) PF01809 - Haemolytic domain (Haemolytic) |
| Potri.016G101700 | | | |
| Potri.016G137900 | | | |
| Potri.016G143100 | AT3G54210 | | Ribosomal protein L17 family protein |
| Potri.017G054000 | AT2G32710 | | Cyclin-dependent kinase inhibitor family protein |
| Potri.017G070100 | AT1G13460 | | Protein phosphatase 2A regulatory B subunit family protein |
| Potri.017G128300 | AT3G03380 | DEG7 | DegP protease 7 |
| Potri.018G057200 | AT2G25850 | PAPS2 | poly(A) polymerase 2 |
| Potri.018G057600 | AT2G19770 | PRF5 | profilin 5 |
| Potri.018G067700 | AT5G56750 | NDL1 | N-MYC downregulated-like 1 |
| Potri.018G070700 | AT5G20060 | | alpha/beta-Hydrolases superfamily protein |
| Potri.018G084900 | AT2G20050 | | protein serine/threonine phosphatases;protein kinases; catalytics; cAMP-dependent protein kinase regulators; ATP binding;protein serine/ threonine phosphatases |
| Potri.018G089900 | AT2G18980 | | Peroxidase superfamily protein |
| Potri.018G096700 | AT4G30340 | DGK7 | diacylglycerol kinase 7 |
| Potri.018G097900 | AT4G16850 | | |
| Potri.018G099400 | AT5G57800 | CER3 | Fatty acid hydroxylase superfamily |
| Potri.018G101300 | AT2G23950 | | Leucine-rich repeat protein kinase family protein |
| Potri.019G041900 | AT3G02890 | | RING/FYVE/PHD zinc finger superfamily protein |
| Potri.019G044400 | AT5G17690 | TFL2 | like heterochromatin protein (LHP1) |
| Potri.019G045800 | AT3G03630 | CS26 | cysteine synthase 26 |
| Potri.019G046400 | AT1G03250 | | (1 of 1) PTHR32019:SF2 - R3H |

| | | | |
|---|---|---|---|
| | | | DOMAIN-CONTAINING PROTEIN 4 |
| Potri.019G047600 | AT3G02930 | | Plant protein of unknown function (DUF827) |
| Potri.019G048100 | AT3G03140 | | Tudor/PWWP/MBT superfamily protein |
| Potri.019G049000 | AT3G10970 | | Haloacid dehalogenase-like hydrolase (HAD) superfamily (HAD) superfamily protein |
| Potri.019G050900 | AT5G16770 | MYB9 | myb domain protein 9 |
| Potri.019G052200 | AT3G03120 | ARFB1C | ADP-ribosylation factor B1C |
| Potri.019G053200 | AT3G02830 | ZFN1 | zinc finger protein 1 |
| Potri.019G054200 | AT3G02750 | | Protein phosphatase 2C family protein |
| Potri.019G066600 | AT1G77350 | | (1 of 1) KOG4615 - Uncharacterized conserved protein |
| Potri.019G081000 | AT3G20230 | | Ribosomal L18p/L5e family protein |
