## Supplemental Table 5 for "Barriers to gene flow play an important role in miantaining reproductive isolation between two closely related *Populus* (Salicaceae) species"

**Table S5.** List of genes located in a region of significantly low genetic differentiation between *P. alba* and *P. adenopoda*.

| Poplar gene | Best Arabidopsis hit | Synonyms | Annotated description |
|---|---|---|---|
| Potri.001G031200 | | | |
| Potri.001G035300 | | | |
| Potri.001G037200 | AT3G14250 | | RING/U-box superfamily protein |
| Potri.001G038000 | AT1G16190 | RAD23A | Rad23 UV excision repair protein family |
| Potri.001G041800 | | | |
| Potri.001G043900 | AT1G15990 | CNGC7 | cyclic nucleotide gated channel 7 |
| Potri.001G044000 | AT1G73060 | LPA3 | Low PSII Accumulation 3 |
| Potri.001G046100 | AT1G80820 | CCR2 | cinnamoyl coa reductase |
| Potri.001G046400 | AT1G80820 | CCR2 | cinnamoyl coa reductase |
| Potri.001G051800 | | | |
| Potri.001G065300 | | | |
| Potri.001G071500 | AT5G47900 | | Protein of unknown function (DUF1624) |
| Potri.001G076300 | | | |
| Potri.001G134500 | | | |
| Potri.001G146800 | AT5G45540 | | Protein of unknown function (DUF594) |
| Potri.001G172600 | AT1G52670 | | Single hybrid motif superfamily protein |
| Potri.001G174500 | AT1G74810 | BOR5 | HCO3- transporter family |
| Potri.001G181200 | AT1G79915 | | Putative methyltransferase family protein |
| Potri.001G189500 | AT1G15210 | ABCG35 | pleiotropic drug resistance 7 |
| Potri.001G208900 | AT5G27010 | | ARM repeat superfamily protein |
| Potri.001G223900 | AT2G44490 | PEN2 | Glycosyl hydrolase superfamily protein |
| Potri.001G236700 | AT3G46010 | ADF1 | actin depolymerizing factor 1 |
| Potri.001G267700 | AT3G63530 | BB | RING/U-box superfamily protein |
| Potri.001G268700 | | | |
| Potri.001G270500 | | | |
| Potri.001G283200 | | | |
| Potri.001G333000 | | | |
| Potri.001G334000 | AT3G01790 | | Ribosomal protein L13 family protein |
| Potri.001G340200 | AT3G01680 | SEOR1 | (1 of 3) PF14576 - Sieve element occlusion N-terminus (SEO_N) |
| Potri.001G340300 | AT3G01680 | SEOR1 | (1 of 3) PF14576 - Sieve element occlusion N-terminus (SEO_N) |
| Potri.001G360800 | AT1G66980 | SNC4 | suppressor of npr1-1 constitutive 4 |
| Potri.001G386300 | AT1G53430 | | Leucine-rich repeat transmembrane protein kinase |
| Potri.001G386700 | AT1G53450 | | |

| | | | |
|---|---|---|---|
| Potri.001G389100 | AT3G11010 | RLP34 | receptor like protein 34 |
| Potri.001G389200 | AT1G05675 | | UDP-Glycosyltransferase superfamily protein |
| Potri.001G395200 | AT2G20562 | | |
| Potri.001G405100 | | | |
| Potri.001G410800 | | | |
| Potri.001G413000 | | | |
| Potri.001G413300 | | | |
| Potri.001G418100 | | | |
| Potri.001G419400 | AT5G54790 | | (1 of 2) PTHR33974:SF2 - GENOMIC DNA, CHROMOSOME 3, P1 CLONE: MSD21 |
| Potri.001G419500 | | | |
| Potri.001G420300 | AT5G54810 | TSB1 | tryptophan synthase beta-subunit 1 |
| Potri.001G420400 | AT3G20550 | DDL | SMAD/FHA domain-containing protein |
| Potri.001G422966 | AT5G54930 | | AT hook motif-containing protein |
| Potri.001G423200 | AT4G26990 | | (1 of 2) PTHR12854//PTHR12854:SF12 - ATAXIN 2-RELATED |
| Potri.001G423400 | AT4G27000 | ATRBP45C | RNA-binding (RRM/RBD/RNP motifs) family protein |
| Potri.001G424600 | AT4G27040 | VPS22 | EAP30/Vps36 family protein |
| Potri.001G425800 | | | |
| Potri.001G429200 | AT2G38130 | ATMAK3 | Acyl-CoA N-acyltransferases (NAT) superfamily protein |
| Potri.001G435400 | | | |
| Potri.001G435900 | AT3G14370 | WAG2 | Protein kinase superfamily protein |
| Potri.001G436600 | AT1G78340 | GSTU22 | glutathione S-transferase TAU 22 |
| Potri.001G440100 | AT5G44500 | | Small nuclear ribonucleoprotein family protein |
| Potri.001G440600 | AT4G20410 | GSNAP | gamma-soluble NSF attachment protein |
| Potri.001G440900 | AT2G34880 | MEE27 | Transcription factor jumonji (jmj) family protein/zinc finger (C5HC2 type) family protein |
| Potri.001G442500 | AT5G44560 | VPS2.2 | SNF7 family protein |
| Potri.001G442900 | AT4G19490 | VPS54 | VPS54 |
| Potri.001G443200 | AT2G35630 | MOR1 | ARM repeat superfamily protein |
| Potri.001G447300 | | | |
| Potri.001G460000 | AT3G44680 | HDA9 | histone deacetylase 9 |
| Potri.001G466500 | | | |
| Potri.001G467000 | AT1G30570 | HERK2 | hercules receptor kinase 2 |
| Potri.001G469600 | AT3G13310 | | Chaperone DnaJ-domain superfamily |

| | | | |
|---|---|---|---|
| | | | protein |
| Potri.002G007800 | AT1G76390 | PUB43 | ARM repeat superfamily protein |
| Potri.002G048666 | | | |
| Potri.002G058300 | AT1G67930 | | Golgi transport complex protein-related |
| Potri.002G201200 | AT4G02220 | | zinc finger (MYND type) family protein / programmed cell death 2 C-terminal domain-containing protein |
| Potri.002G201300 | AT2G47510 | FUM1 | fumarase 1 |
| Potri.002G201400 | AT1G33390 | FAS4 | RNA helicase family protein |
| Potri.002G222700 | AT5G35735 | | Auxin-responsive family protein |
| Potri.002G238500 | | | |
| Potri.002G255700 | AT5G40120 | AGL76 | AGAMOUS-like 76 |
| Potri.003G000900 | AT5G49880 | MAD1 | mitotic checkpoint family protein |
| Potri.003G003100 | AT4G18530 | | Protein of unknown function (DUF707) |
| Potri.003G004100 | AT1G09630 | RAB11c | RAB GTPase 11C |
| Potri.003G004700 | AT1G09640 | | Translation elongation factor EF1B, gamma chain |
| Potri.003G005500 | AT1G09660 | | RNA-binding KH domain-containing protein |
| Potri.003G006900 | AT1G58007 | | |
| Potri.003G015000 | | | |
| Potri.003G015700 | AT1G22940 | TH1 | thiamin biosynthesis protein, putative |
| Potri.003G015800 | AT1G60930 | RECQ4B | RECQ helicase L4B |
| Potri.003G018500 | AT1G22930 | | T-complex protein 11 |
| Potri.003G020800 | AT1G33270 | | Acyl transferase/acyl hydrolase/lysophos-pholipase superfamily protein |
| Potri.003G023500 | | | |
| Potri.003G023700 | AT5G09390 | | CD2-binding protein-related |
| Potri.003G024000 | AT4G33520 | PAA1 | P-type ATP-ase 1 |
| Potri.003G024100 | AT3G02645 | | Plant protein of unknown function (DUF247) |
| Potri.003G024200 | AT1G60500 | DRP4C | Dynamin related protein 4C |
| Potri.003G024600 | AT1G60500 | DRP4C | Dynamin related protein 4C |
| Potri.003G024800 | AT1G60500 | DRP4C | Dynamin related protein 4C |
| Potri.003G024900 | AT1G60500 | DRP4C | Dynamin related protein 4C |
| Potri.003G025566 | | | |
| Potri.003G026500 | AT1G53450 | | |
| Potri.003G034700 | AT3G21250 | ABCC8 | multidrug resistance-associated protein 6 |
| Potri.003G042300 | | | |
| Potri.003G059900 | AT3G15630 | | |
| Potri.003G060500 | AT1G15440 | PWP2 | periodic tryptophan protein 2 |
| Potri.003G074800 | AT1G70190 | | Ribosomal protein L7/L12, oligomerisation; |

| | | | |
|---|---|---|---|
| | | | Ribosomal protein L7/L12, C-terminal/ adaptor protein ClpS-like |
| Potri.003G095400 | AT5G46870 | | RNA-binding (RRM/RBD/RNP motifs) family protein |
| Potri.003G150000 | | | |
| Potri.003G185300 | AT1G79700 | WRI4 | Integrase-type DNA-binding superfamily protein |
| Potri.003G186100 | AT1G16260 | | Wall-associated kinase family protein |
| Potri.003G186200 | | | |
| Potri.003G188700 | AT5G04940 | SUVH1 | SU(VAR)3-9 homolog 1 |
| Potri.003G189100 | | | |
| Potri.003G189700 | AT3G11440 | MYB65 | myb domain protein 65 |
| Potri.003G190300 | | | |
| Potri.003G190400 | | | |
| Potri.003G190900 | AT4G02550 | | (1 of 14) PF12776 - Myb/SANT-like DNA -binding domain (Myb_DNA-bind_3) |
| Potri.003G201400 | AT3G54570 | | Plant calmodulin-binding protein-related |
| Potri.003G201700 | | | |
| Potri.003G201900 | | | |
| Potri.003G202100 | | | |
| Potri.003G205900 | | | |
| Potri.003G206000 | AT5G12960 | | Putative glycosyl hydrolase of unknown function (DUF1680) |
| Potri.003G207000 | AT3G20820 | | Leucine-rich repeat (LRR) family protein |
| Potri.003G210100 | AT5G10400 | | Histone superfamily protein |
| Potri.003G211200 | AT3G57930 | | (1 of 2) PTHR34055:SF4 - EXPRESSED PROTEIN |
| Potri.003G212000 | AT5G60900 | RLK1 | receptor-like protein kinase 1 |
| Potri.003G212500 | AT2G42150 | | DNA-binding bromodomain-containing protein |
| Potri.003G212600 | AT3G20760 | | Nse4, component of Smc5/6 DNA repair complex |
| Potri.003G213700 | | | |
| Potri.004G006000 | AT4G04340 | | ERD (early-responsive to dehydration stress) family protein |
| Potri.004G007000 | AT1G11925 | | Stigma-specific Stig1 family protein |
| Potri.004G007200 | AT1G11925 | | Stigma-specific Stig1 family protein |
| Potri.004G008700 | AT4G04470 | PMP22 | Peroxisomal membrane 22 kDa (Mpv17/ PMP22) family protein |
| Potri.004G009500 | AT4G22030 | | F-box family protein with a domain of unknown function (DUF295) |

| | | | |
|---|---|---|---|
| Potri.004G010900 | | | |
| Potri.004G011000 | AT3G53310 | | AP2/B3-like transcriptional factor family protein |
| Potri.004G014450 | AT4G18250 | | receptor serine/threonine kinase, putative |
| Potri.004G014700 | AT4G18250 | | receptor serine/threonine kinase, putative |
| Potri.004G020000 | | | |
| Potri.004G024101 | | | |
| Potri.004G025800 | AT4G23270 | CRK19 | cysteine-rich RLK (RECEPTOR-like protein kinase) 19 |
| Potri.004G025900 | AT4G23270 | CRK19 | cysteine-rich RLK (RECEPTOR-like protein kinase) 19 |
| Potri.004G027400 | AT1G11350 | SD1-13 | S-domain-1 13 |
| Potri.004G028800 | AT4G21350 | PUB8 | plant U-box 8 |
| Potri.004G034700 | | | |
| Potri.004G034800 | | | |
| Potri.004G040400 | AT4G04920 | SFR6 | sensitive to freezing 6 |
| Potri.004G063300 | AT2G34480 | | Ribosomal protein L18ae/LX family protein |
| Potri.004G071000 | AT3G16520 | UGT88A1 | UDP-glucosyl transferase 88A1 |
| Potri.004G091600 | AT5G17460 | | |
| Potri.004G093800 | AT3G29590 | AT5MAT | HXXXD-type acyl-transferase family protein |
| Potri.004G097100 | AT1G66880 | | Protein kinase superfamily protein |
| Potri.004G101900 | | | |
| Potri.004G135600 | AT2G30900 | TBL43 | TRICHOME BIREFRINGENCE-LIKE 43 |
| Potri.004G157300 | AT4G34560 | | (1 of 11) PF14364 - Domain of unknown function (DUF4408) |
| Potri.004G171300 | AT4G34880 | | Amidase family protein |
| Potri.004G189400 | AT5G22750 | RAD5 | DNA/RNA helicase protein |
| Potri.004G191400 | | | |
| Potri.004G191500 | | | |
| Potri.004G193300 | AT5G20930 | TSL | Protein kinase superfamily protein |
| Potri.004G193800 | AT5G12230 | MED19A | (1 of 2) PTHR22536 - LUNG CANCER MET-ASTASIS-RELATED LCMR1 PROTEIN |
| Potri.004G212800 | AT1G08080 | ACA7 | alpha carbonic anhydrase 7 |
| Potri.004G225350 | | | |
| Potri.004G231200 | | | |
| Potri.004G231500 | | | |
| Potri.004G231600 | | | |
| Potri.005G003700 | AT4G08570 | | Heavy metal transport/detoxification superfamily protein |

| | | | |
|---|---|---|---|
| Potri.005G005500 | | | |
| Potri.005G007300 | AT1G10630 | ARFA1F | ADP-ribosylation factor A1F |
| Potri.005G009900 | | | |
| Potri.005G010133 | AT1G08060 | MOM | ATP-dependent helicase family protein |
| Potri.005G010266 | AT1G09340 | CRB | chloroplast RNA binding |
| Potri.005G010600 | AT1G56590 | ZIP4 | Clathrin adaptor complexes medium subunit family protein |
| Potri.005G010800 | AT1G56560 | A/N-InvA | Plant neutral invertase family protein |
| Potri.005G011100 | AT1G09320 | | agenet domain-containing protein |
| Potri.005G011300 | AT1G09280 | | (1 of 1) PTHR22778:SF0 - OVARIAN CAN-CER ASSOCIATED GENE 2 PROTEIN |
| Potri.005G011800 | AT3G05810 | | (1 of 2) PF09597 - IGR protein motif (IGR) |
| Potri.005G012400 | AT1G09250 | | basic helix-loop-helix (bHLH) DNA-binding superfamily protein |
| Potri.005G012700 | | | |
| Potri.005G012900 | AT5G26820 | IREG3 | iron-regulated protein 3 |
| Potri.005G013800 | AT1G54470 | RPP27 | RNI-like superfamily protein |
| Potri.005G014100 | AT1G56440 | TPR5 | Tetratricopeptide repeat (TPR)-like superfamily protein |
| Potri.005G014200 | | | |
| Potri.005G014700 | | | |
| Potri.005G017600 | AT5G26860 | LON1 | lon protease 1 |
| Potri.005G030271 | AT5G27550 | | P-loop containing nucleoside triphosphate hydrolases superfamily protein |
| Potri.005G030413 | AT4G04110 | | Toll-Interleukin-Resistance (TIR) domain family protein |
| Potri.005G033400 | AT1G54960 | NP2 | NPK1-related protein kinase 2 |
| Potri.005G037000 | AT3G05160 | | Major facilitator superfamily protein |
| Potri.005G038100 | AT5G27380 | GSH2 | glutathione synthetase 2 |
| Potri.005G039100 | AT3G05190 | | D-aminoacid aminotransferase-like PLP-dependent enzymes superfamily protein |
| Potri.005G039500 | AT2G29370 | | NAD(P)-binding Rossmann-fold superfamily protein |
| Potri.005G042700 | AT2G32910 | | DCD (Development and Cell Death) domain protein |
| Potri.005G043100 | AT2G32910 | | DCD (Development and Cell Death) domain protein |
| Potri.005G043200 | AT3G04980 | | DNAJ heat shock N-terminal domain-containing protein |
| Potri.005G044000 | AT5G27230 | | Frigida-like protein |

| | | | |
|---|---|---|---|
| Potri.005G044600 | AT4G08630 | | |
| Potri.005G050300 | AT1G63150 | | Tetratricopeptide repeat (TPR)-like superfamily protein |
| Potri.005G079900 | AT5G10080 | | Eukaryotic aspartyl protease family protein |
| Potri.005G115601 | AT1G07660 | | Histone superfamily protein |
| Potri.005G127100 | AT4G36810 | GGPS1 | geranylgeranyl pyrophosphate synthase 1 |
| Potri.005G196400 | AT4G09830 | | Uncharacterised conserved protein UCP009193 |
| Potri.006G010500 | | | |
| Potri.006G011200 | | | |
| Potri.006G011300 | AT4G21060 | | Galactosyltransferase family protein |
| Potri.006G012900 | | | |
| Potri.006G013200 | AT4G22390 | | F-box associated ubiquitination effector family protein |
| Potri.006G013500 | | | |
| Potri.006G014232 | | | |
| Potri.006G028000 | | | |
| Potri.006G060300 | | | |
| Potri.006G062050 | | | |
| Potri.006G062500 | AT5G20550 | | 2-oxoglutarate (2OG) and Fe(II)-dependent oxygenase superfamily protein |
| Potri.006G063300 | AT5G20410 | MGD2 | monogalactosyldiacylglycerol synthase 2 |
| Potri.006G063500 | AT5G20300 | Toc90 | Avirulence induced gene (AIG1) family protein |
| Potri.006G119800 | AT3G53130 | LUT1 | Cytochrome P450 superfamily protein |
| Potri.006G124400 | AT3G46100 | HRS1 | Histidyl-tRNA synthetase 1 |
| Potri.006G149500 | AT4G29490 | | Metallopeptidase M24 family protein |
| Potri.006G180300 | AT4G30840 | | Transducin/WD40 repeat-like superfamily protein |
| Potri.006G187000 | | | |
| Potri.006G241800 | AT4G32720 | La1 | La protein 1 |
| Potri.006G268700 | AT2G29120 | GLR2.7 | glutamate receptor 2.7 |
| Potri.006G268900 | AT2G29120 | GLR2.7 | glutamate receptor 2.7 |
| Potri.006G272600 | | | |
| Potri.006G272800 | AT2G24580 | | FAD-dependent oxidoreductase family protein |
| Potri.006G272900 | AT4G31540 | EXO70G1 | exocyst subunit exo70 family protein G1 |
| Potri.006G275600 | AT4G31390 | ACDO1 | Protein kinase superfamily protein |
| Potri.006G275700 | AT2G24490 | RPA2 | replicon protein A2 |
| Potri.007G088500 | AT4G39810 | | Polynucleotidyl transferase, ribonuclease H-like superfamily protein |

| | | | |
|---|---|---|---|
| Potri.007G111500 | | | |
| Potri.007G117200 | AT2G43840 | UGT74F1 | UDP-glycosyltransferase 74 F1 |
| Potri.007G123000 | | | |
| Potri.007G124700 | AT2G43250 | | |
| Potri.007G125450 | | | |
| Potri.007G126100 | | | |
| Potri.007G133400 | AT3G59580 | | Plant regulator RWP-RK family protein |
| Potri.007G136000 | | | |
| Potri.007G141900 | | | |
| Potri.007G142100 | AT2G43850 | | Integrin-linked protein kinase family |
| Potri.008G003000 | AT5G23390 | | Plant protein of unknown function (DUF639) |
| Potri.008G004400 | AT3G20260 | | Protein of unknown function (DUF1666) |
| Potri.008G005000 | AT1G78900 | VHA-A | vacuolar ATP synthase subunit A |
| Potri.008G033500 | | | |
| Potri.008G143000 | | | |
| Potri.008G185500 | AT3G26430 | | GDSL-like Lipase/Acylhydrolase superfamily protein |
| Potri.008G202500 | AT5G19060 | | |
| Potri.008G220700 | AT3G06868 | | |
| Potri.009G036300 | | | |
| Potri.009G141700 | AT3G12500 | HCHIB | basic chitinase |
| Potri.009G152800 | | | |
| Potri.009G153300 | AT3G43610 | | Spc97 / Spc98 family of spindle pole body (SBP) component |
| Potri.009G153600 | AT1G04580 | AO4 | aldehyde oxidase 4 |
| Potri.009G153900 | AT5G20950 | | Glycosyl hydrolase family protein |
| Potri.009G154900 | | | |
| Potri.010G000400 | | | |
| Potri.010G017800 | | | |
| Potri.010G018300 | | | |
| Potri.010G020200 | | | |
| Potri.010G025800 | | | |
| Potri.010G044400 | AT1G10740 | | alpha/beta-Hydrolases superfamily protein |
| Potri.010G044500 | AT1G23320 | TAR1 | tryptophan aminotransferase related 1 |
| Potri.010G044700 | AT1G60780 | HAP13 | Clathrin adaptor complexes medium subunit family protein |
| Potri.010G044900 | AT1G70570 | | anthranilate phosphoribosyltransferase, putative |
| Potri.010G050300 | AT3G01680 | | (1 of 3) PF14576 - Sieve element occlusion N-terminus (SEO_N) |

| | | | |
|---|---|---|---|
| Potri.010G053200 | AT3G01710 | | TPX2 (targeting protein for Xklp2) protein family |
| Potri.010G070800 | AT3G23280 | XBAT35 | XB3 ortholog 5 in Arabidopsis thaliana |
| Potri.010G119700 | AT1G15060 | | Uncharacterised conserved protein UCP031088, alpha/beta hydrolase |
| Potri.010G130800 | AT3G25690 | CHUP1 | Hydroxyproline-rich glycoprotein family protein |
| Potri.010G155150 | | | |
| Potri.010G155400 | AT4G17270 | | Mo25 family protein |
| Potri.010G195300 | AT3G55710 | | UDP-Glycosyltransferase superfamily protein |
| Potri.010G196800 | | | |
| Potri.010G253200 | AT1G50430 | DWF5 | Ergosterol biosynthesis ERG4/ERG24 family |
| Potri.011G005000 | AT4G22150 | PUX3 | plant UBX domain-containing protein 3 |
| Potri.011G008100 | AT4G22090 | | Pectin lyase-like superfamily protein |
| Potri.011G011300 | AT4G22150 | | plant UBX domain-containing protein 3 |
| Potri.011G015500 | AT4G22190 | | |
| Potri.011G019500 | | | |
| Potri.011G021400 | | | |
| Potri.011G028400 | AT4G23270 | | cysteine-rich RLK (RECEPTOR-like protein kinase) 19 |
| Potri.011G028500 | | | |
| Potri.011G028700 | | | |
| Potri.011G029300 | AT4G21400 | CRK28 | cysteine-rich RLK (RECEPTOR-like protein kinase) 28 |
| Potri.011G029500 | | | |
| Potri.011G030000 | AT4G21400 | CRK28 | cysteine-rich RLK (RECEPTOR-like protein kinase) 28 |
| Potri.011G030200 | | | |
| Potri.011G030400 | | | |
| Potri.011G035700 | | | |
| Potri.011G037800 | AT1G11350 | | S-domain-1 13 |
| Potri.011G038401 | AT1G11350 | | S-domain-1 13 |
| Potri.011G038901 | | | |
| Potri.011G039100 | AT1G11350 | | S-domain-1 13 |
| Potri.011G039400 | AT1G11350 | | S-domain-1 13 |
| Potri.011G039500 | | | |
| Potri.011G040000 | AT1G61620 | | phosphoinositide binding |
| Potri.011G045300 | AT1G11170 | | Protein of unknown function (DUF707) |
| Potri.011G062500 | AT1G29030 | | Apoptosis inhibitory protein 5 (API5) |

| | | | |
|---|---|---|---|
| Potri.011G062750 | | | |
| Potri.011G062900 | | | |
| Potri.011G064800 | AT1G29070 | | Ribosomal protein L34 |
| Potri.011G072566 | AT1G29750 | RKF1 | receptor-like kinase in flowers 1 |
| Potri.011G080600 | AT4G18270 | TRANS11 | translocase 11 |
| Potri.011G082100 | AT5G46150 | | LEM3 (ligand-effect modulator 3) family protein /CDC50 family protein |
| Potri.011G095700 | AT5G27010 | | ARM repeat superfamily protein |
| Potri.011G096400 | | | |
| Potri.011G104900 | | | |
| Potri.011G105500 | AT1G78610 | MSL6 | mechanosensitive channel of small conductance-like 6 |
| Potri.011G108500 | AT1G53330 | | Pentatricopeptide repeat (PPR) superfamily protein |
| Potri.011G117600 | | | |
| Potri.011G117700 | | | |
| Potri.011G118000 | AT5G53860 | emb2737 | embryo defective 2737 |
| Potri.011G137300 | AT5G54860 | | Major facilitator superfamily protein |
| Potri.011G138900 | | | |
| Potri.011G163000 | AT5G44410 | | FAD-binding Berberine family protein |
| Potri.011G165700 | | | |
| Potri.012G002800 | | | |
| Potri.012G004400 | AT3G49590 | ATG13 | Autophagy-related protein 13 |
| Potri.012G006100 | AT5G24520 | TTG1 | Transducin/WD40 repeat-like superfamily protein |
| Potri.012G007400 | AT5G54140 | ILL3 | IAA-leucine-resistant (ILR1)-like 3 |
| Potri.012G007600 | AT5G54130 | | Calcium-binding endonuclease/exonuclease/ phosphatase family |
| Potri.012G007700 | | | |
| Potri.012G011000 | | | |
| Potri.012G012700 | AT5G24170 | | Got1/Sft2-like vescicle transport protein family |
| Potri.012G015800 | AT4G22000 | | |
| Potri.012G016600 | AT5G53020 | | Ribonuclease P protein subunit P38-related |
| Potri.012G017400 | | | |
| Potri.012G018300 | AT5G24330 | ATXR6 | ARABIDOPSIS TRITHORAX-RELATED PROTEIN 6 |
| Potri.012G018800 | | | |
| Potri.012G019500 | AT4G27730 | OPT6 | oligopeptide transporter 1 |
| Potri.012G020200 | AT1G55580 | LAS | GRAS family transcription factor |
| Potri.012G020700 | AT4G27700 | | Rhodanese/Cell cycle control phosphatase |

| | | | |
|---|---|---|---|
| | | | superfamily protein |
| Potri.012G020800 | | | |
| Potri.012G024100 | | | |
| Potri.012G024300 | AT3G52590 | UBQ1 | ubiquitin extension protein 1 |
| Potri.012G024500 | AT5G24390 | | Ypt/Rab-GAP domain of gyp1p superfamily protein |
| Potri.012G031000 | AT5G24130 | | |
| Potri.012G031100 | AT5G24120 | SIGE | sigma factor E |
| Potri.012G039700 | AT1G73450 | | Protein kinase superfamily protein |
| Potri.012G044400 | | | |
| Potri.012G048200 | AT1G73630 | | EF hand calcium-binding protein family |
| Potri.012G054200 | AT3G18270 | CYP77A5P | cytochrome P450, family 77, subfamily A, polypeptide 5 pseudogene |
| Potri.012G055000 | AT1G48760 | delta-ADR | delta-adaptin |
| Potri.012G055700 | AT1G18400 | BEE1 | BR enhanced expression 1 |
| Potri.012G056700 | | | |
| Potri.012G071100 | AT5G62710 | | Leucine-rich repeat protein kinase family protein |
| Potri.012G089700 | AT4G24740 | FC2 | FUS3-complementing gene 2 |
| Potri.012G123550 | AT1G79210 | | N-terminal nucleophile aminohydrolases (Ntn hydrolases) superfamily protein |
| Potri.012G123700 | AT2G35120 | | Single hybrid motif superfamily protein |
| Potri.012G124500 | AT1G31170 | SRX | sulfiredoxin |
| Potri.012G125500 | AT5G05010 | | clathrin adaptor complexes medium subunit family protein |
| Potri.013G037700 | AT3G04870 | ZDS | zeta-carotene desaturase |
| Potri.013G037900 | | | |
| Potri.013G038700 | AT5G28220 | | Protein prenylyltransferase superfamily protein |
| Potri.013G051200 | AT5G18410 | PIR121 | transcription activators |
| Potri.013G060000 | | | |
| Potri.013G072300 | AT5G17620 | 7-Aug | (1 of 1) PTHR14352:SF2 - HAUS AUGMIN-LIKECOMPLEX SUBUNIT 7 |
| Potri.013G098000 | | | |
| Potri.013G102900 | | | |
| Potri.013G114500 | AT1G22700 | | Tetratricopeptide repeat (TPR)-like superfamily protein |
| Potri.013G121000 | AT2G19130 | | S-locus lectin protein kinase family protein |
| Potri.013G124600 | AT1G60500 | | Dynamin related protein 4C |
| Potri.013G126300 | AT1G73180 | | Eukaryotic translation initiation factor eIF2A family protein |

| | | | |
|---|---|---|---|
| Potri.013G126701 | AT5G60030 | | |
| Potri.013G128100 | | | |
| Potri.013G128500 | AT3G59770 | SAC9 | sacI homology domain-containing protein / WW domain-containing protein |
| Potri.013G128600 | AT3G25520 | ATL5 | ribosomal protein L5 |
| Potri.013G128900 | AT1G17680 | | tetratricopeptide repeat (TPR)-containing protein |
| Potri.013G129600 | | | |
| Potri.013G131100 | AT4G03560 | TPC1 | two-pore channel 1 |
| Potri.013G134200 | AT2G20740 | | Tetraspanin family protein |
| Potri.013G134700 | AT2G20770 | GCL2 | GCR2-like 2 |
| Potri.013G136100 | AT2G20875 | EPF1 | epidermal patterning factor 1 |
| Potri.013G143800 | | | |
| Potri.013G144300 | AT4G03400 | DFL2 | Auxin-responsive GH3 family protein |
| Potri.013G145400 | AT4G28190 | ULT1 | Developmental regulator, ULTRAPETALA |
| Potri.013G147600 | AT1G12470 | | zinc ion binding |
| Potri.013G149300 | | | |
| Potri.013G151432 | | | |
| Potri.013G152400 | AT4G03210 | XTH9 | xyloglucan endotransglucosylase/ hydrolase 9 |
| Potri.013G157250 | | | |
| Potri.014G004500 | AT4G37950 | | Rhamnogalacturonate lyase family protein |
| Potri.014G004900 | AT3G49850 | TRB3 | telomere repeat binding factor 3 |
| Potri.014G007000 | | | |
| Potri.014G010000 | AT2G22640 | BRK1 | BRICK1, putative |
| Potri.014G010100 | AT5G10540 | | Zincin-like metalloproteases family protein |
| Potri.014G012100 | AT5G65840 | | Thioredoxin superfamily protein |
| Potri.014G037400 | | | |
| Potri.014G037600 | | | |
| Potri.014G053700 | | | |
| Potri.015G001000 | AT1G04635 | EMB1687 | ribonuclease P family protein / Rpp14 family |
| Potri.015G014850 | AT3G53310 | | AP2/B3-like transcriptional factor family protein |
| Potri.015G016000 | AT3G53310 | | AP2/B3-like transcriptional factor family protein |
| Potri.015G016450 | AT3G53310 | | AP2/B3-like transcriptional factor family protein |
| Potri.015G018700 | AT3G49170 | EMB2261 | Tetratricopeptide repeat (TPR)-like superfamily protein |
| Potri.015G022400 | AT5G41480 | GLA1 | Folylpolyglutamate synthetase family protein |
| Potri.015G022700 | AT4G27940 | MTM1 | manganese tracking factor for mitochondrial |

| | | | |
|---|---|---|---|
| | | | SOD2 |
| Potri.015G023900 | | | |
| Potri.015G024200 | AT5G24090 | CHIA | chitinase A |
| Potri.015G024400 | AT3G26560 | | ATP-dependent RNA helicase, putative |
| Potri.015G034000 | AT3G17690 | CNGC19 | cyclic nucleotide gated channel 19 |
| Potri.015G064300 | AT1G18850 | | |
| Potri.015G064900 | | | |
| Potri.015G079100 | | | |
| Potri.015G082500 | AT5G63110 | HDA6 | histone deacetylase 6 |
| Potri.015G102200 | AT3G48770 | | DNA binding;ATP binding |
| Potri.015G102700 | AT3G48770 | | DNA binding;ATP binding |
| Potri.015G103000 | AT3G48770 | | DNA binding;ATP binding |
| Potri.015G112500 | AT5G45170 | | Haloacid dehalogenase-like hydrolase (HAD) superfamily protein |
| Potri.015G112600 | | | |
| Potri.015G118200 | AT1G31240 | | Bromodomain transcription factor |
| Potri.015G118500 | AT1G10180 | | (1 of 1) PTHR21426:SF2 - EXOCYST COMPLEX COMPONENT EXO84C |
| Potri.015G121400 | AT2G35155 | | Trypsin family protein |
| Potri.015G122300 | | | |
| Potri.015G122400 | AT1G79210 | | N-terminal nucleophile aminohydrolases (Ntn hydrolases) superfamily protein |
| Potri.015G122700 | | | |
| Potri.015G125950 | | | |
| Potri.016G009300 | | | |
| Potri.016G009400 | | | |
| Potri.016G010600 | AT2G35740 | INT3 | nositol transporter 3 |
| Potri.016G010900 | AT4G17280 | | Auxin-responsive family protein |
| Potri.016G011100 | AT4G17270 | | Mo25 family protein |
| Potri.016G012700 | | | |
| Potri.016G014300 | | | |
| Potri.016G015800 | | | |
| Potri.016G016700 | AT3G21800 | UGT71B8 | UDP-glucosyl transferase 71B8 |
| Potri.016G022600 | | | |
| Potri.016G027900 | | | |
| Potri.016G028200 | | | |
| Potri.016G047600 | | | |
| Potri.016G055100 | AT2G41830 | | Uncharacterized protein |
| Potri.016G062600 | AT3G11964 | | RNA binding;RNA binding |
| Potri.016G092600 | | | |
| Potri.016G093000 | AT5G03370 | | acylphosphatase family |

| | | | |
|---|---|---|---|
| Potri.016G102700 | | | |
| Potri.016G105400 | | | |
| Potri.016G110700 | AT5G01290 | | mRNA capping enzyme family protein |
| Potri.016G120100 | AT5G01320 | | Thiamine pyrophosphate dependent pyruvate decarboxylase family protein |
| Potri.016G120600 | | | |
| Potri.016G126400 | AT2G38360 | PRA1.B4 | prenylated RAB acceptor 1.B4 |
| Potri.016G127500 | AT3G08800 | SIEL | ARM repeat superfamily protein |
| Potri.017G000800 | | | |
| Potri.017G002300 | AT5G16960 | | Zinc-binding dehydrogenase family protein |
| Potri.017G005400 | | | |
| Potri.017G006700 | AT2G43890 | | Pectin lyase-like superfamily protein |
| Potri.017G009500 | | | |
| Potri.017G009600 | | | |
| Potri.017G010800 | | | |
| Potri.017G012600 | AT2G20650 | | RING/U-box superfamily protein |
| Potri.017G014300 | AT4G28430 | | Reticulon family protein |
| Potri.017G014600 | AT5G44200 | CBP20 | CAP-binding protein 20 |
| Potri.017G015600 | AT3G14470 | | NB-ARC domain-containing disease resistance protein |
| Potri.017G023300 | | | |
| Potri.017G029200 | AT4G22720 | | Actin-like ATPase superfamily protein |
| Potri.017G029400 | AT2G43280 | | Far-red impaired responsive (FAR1) family protein |
| Potri.017G031800 | AT2G47680 | | zinc finger (CCCH type) helicase family protein |
| Potri.017G033300 | AT2G43360 | BIO2 | Radical SAM superfamily protein |
| Potri.017G035700 | AT2G43250 | | |
| Potri.017G035800 | AT2G43240 | | Nucleotide-sugar transporter family protein |
| Potri.017G036200 | AT1G54380 | | spliceosome protein-related |
| Potri.017G036600 | AT2G43210 | | Ubiquitin-like superfamily protein |
| Potri.017G037701 | AT3G05870 | APC11 | anaphase-promoting complex/cyclosome 11 |
| Potri.017G043200 | AT4G33625 | | (1 of 1) PF08507 - COPI associated protein (COPI_assoc) |
| Potri.017G043600 | AT1G29490 | | SAUR-like auxin-responsive protein family |
| Potri.017G047700 | | | |
| Potri.017G048600 | AT4G05440 | EDA35 | temperature sensing protein-related |
| Potri.017G049000 | | | |
| Potri.017G050800 | AT5G64270 | | splicing factor, putative |
| Potri.017G062900 | AT3G02645 | | Plant protein of unknown function (DUF247) |

| | | | |
|---|---|---|---|
| Potri.017G072100 | AT3G27700 | | zinc finger (CCCH-type) family protein / RNA recognition motif (RRM)-containing protein |
| Potri.017G080100 | AT3G01260 | | Galactose mutarotase-like superfamily protein |
| Potri.017G081400 | AT3G01210 | | RNA-binding (RRM/RBD/RNP motifs) family protein |
| Potri.017G096000 | | | |
| Potri.017G098500 | AT5G15640 | | Mitochondrial substrate carrier family protein |
| Potri.017G099200 | AT5G15810 | | N2,N2-dimethylguanosine tRNA methyltransferase |
| Potri.017G099800 | | | |
| Potri.017G102500 | AT3G30380 | | alpha/beta-Hydrolases superfamily protein |
| Potri.017G102700 | AT5G64130 | | cAMP-regulated phosphoprotein 19-related protein |
| Potri.017G115900 | AT5G20480 | EFR | EF-TU receptor |
| Potri.017G116700 | AT5G16180 | CRS1 | ortholog of maize chloroplast splicing factor CRS1 |
| Potri.017G119700 | | | |
| Potri.017G123900 | AT1G66510 | | AAR2 protein family |
| Potri.017G131100 | AT1G66200 | GSR2 | glutamine synthase clone F11 |
| Potri.017G133300 | | | |
| Potri.017G136950 | | | |
| Potri.017G138800 | AT3G03190 | GSTF11 | glutathione S-transferase F11 |
| Potri.017G140200 | AT1G65810 | | P-loop containing nucleoside triphosphate hydrolases superfamily protein |
| Potri.017G140400 | | | |
| Potri.017G141400 | | | |
| Potri.017G142400 | AT5G52510 | SCL8 | SCARECROW-like 8 |
| Potri.017G144700 | AT3G03250 | UGP1 | UDP-GLUCOSE PYROPHOSPHOR-YLASE 1 |
| Potri.017G144800 | AT5G17300 | RVE1 | Homeodomain-like superfamily protein |
| Potri.017G145100 | | | |
| Potri.017G145550 | | | |
| Potri.017G151600 | AT1G48370 | YSL8 | YELLOW STRIPE like 8 |
| Potri.018G000101 | | | |
| Potri.018G001000 | AT4G31150 | | endonuclease V family protein |
| Potri.018G001301 | | | |
| Potri.018G011800 | AT2G29120 | | glutamate receptor 2.7 |
| Potri.018G012100 | AT2G29120 | | glutamate receptor 2.7 |

| | | | |
|---|---|---|---|
| Potri.018G012600 | AT2G29120 | | glutamate receptor 2.7 |
| Potri.018G012900 | AT2G29120 | | glutamate receptor 2.7 |
| Potri.018G014600 | | | |
| Potri.018G016600 | AT4G31780 | MGD1 | monogalactosyl diacylglycerol synthase 1 |
| Potri.018G111700 | | | |
| Potri.018G111800 | | | |
| Potri.018G111900 | | | |
| Potri.018G121600 | AT5G20380 | PHT4;5 | phosphate transporter 4;5 |
| Potri.018G121800 | AT5G20550 | | 2-oxoglutarate (2OG) and Fe(II)-dependent oxygenase superfamily protein |
| Potri.018G130200 | | | |
| Potri.018G133500 | AT2G19350 | | Eukaryotic protein of unknown function (DUF872) |
| Potri.018G135600 | | | |
| Potri.018G137500 | AT5G23575 | | Transmembrane CLPTM1 family protein |
| Potri.018G138200 | AT5G26360 | | TCP-1/cpn60 chaperonin family protein |
| Potri.018G140900 | AT4G29920 | | Double Clp-N motif-containing P-loop nucleoside triphosphate hydrolases superfamily protein |
| Potri.018G141900 | AT5G57160 | ATLIG4 | DNA ligase IV |
| Potri.018G142400 | AT4G29940 | PRHA | pathogenesis related homeodomain protein A |
| Potri.018G143100 | AT2G19240 | | Ypt/Rab-GAP domain of gyp1p superfamily protein |
| Potri.018G143400 | AT4G30020 | | PA-domain containing subtilase family protein |
| Potri.018G143800 | AT5G57250 | | Pentatricopeptide repeat (PPR) superfamily protein |
| Potri.018G145600 | | | |
| Potri.018G145700 | | | |
| Potri.018G145800 | | | |
| Potri.018G146001 | | | |
| Potri.018G146200 | AT5G09520 | PELPK2 | hydroxyproline-rich glycoprotein family protein |
| Potri.018G146300 | AT5G35360 | CAC2 | acetyl Co-enzyme a carboxylase biotin carboxylase subunit |
| Potri.018G146700 | AT5G35330 | MBD02 | methyl-CPG-binding domain protein 02 |
| Potri.018G146900 | AT3G42630 | | Pentatricopeptide repeat (PPR) superfamily protein |
| Potri.018G147200 | | | |
| Potri.018G149300 | AT5G36110 | CYP716A1 | cytochrome P450, family 716, subfamily A, |

| | | | |
|---|---|---|---|
| | | | polypeptide 1 |
| Potri.018G152200 | AT5G08370 | AGAL2 | alpha-galactosidase 2 |
| Potri.019G001500 | | | |
| Potri.019G001800 | AT5G63710 | | Leucine-rich repeat protein kinase family protein |
| Potri.019G002100 | AT2G03200 | | Eukaryotic aspartyl protease family protein |
| Potri.019G002500 | | | |
| Potri.019G002900 | AT3G03480 | CHAT | acetyl CoA:(Z)-3-hexen-1-ol acetyltrans-ferase |
| Potri.019G003100 | AT1G05520 | | Sec23/Sec24 protein transport family protein |
| Potri.019G003400 | | | |
| Potri.019G003900 | AT5G09850 | | Transcription elongation factor (TFIIS) family protein |
| Potri.019G004000 | AT5G64710 | | Putative endonuclease or glycosyl hydrolase |
| Potri.019G004100 | AT5G64700 | UMAMIT21 | nodulin MtN21 /EamA-like transporter family protein |
| Potri.019G004300 | AT4G28520 | CRU3 | cruciferin 3 |
| Potri.019G004500 | AT4G28520 | CRU3 | cruciferin 3 |
| Potri.019G007600 | | | |
| Potri.019G009700 | | | |
| Potri.019G009800 | | | |
| Potri.019G010400 | AT5G09810 | ACT7 | actin 7 |
| Potri.019G010700 | AT3G18770 | | Autophagy-related protein 13 |
| Potri.019G011400 | | | |
| Potri.019G012200 | AT1G08800 | | Protein of unknown function, DUF593 |
| Potri.019G013800 | AT1G54560 | XIE | Myosin family protein with Dil domain |
| Potri.019G014100 | AT2G14045 | | (1 of 1) PTHR13168 - ASSOCIATE OF C-MYC AMY-1 |
| Potri.019G018200 | | | |
| Potri.019G018800 | | | |
| Potri.019G020000 | AT3G04460 | PEX12 | peroxin-12 |
| Potri.019G020800 | | | |
| Potri.019G021500 | AT1G08600 | ATRX | P-loop containing nucleoside triphosphate hydrolases superfamily protein |
| Potri.019G030300 | AT2G38430 | | |
| Potri.019G039900 | AT4G08580 | CEL5 | microfibrillar-associated protein-related |
| Potri.019G053400 | AT5G36290 | | Uncharacterized protein family (UPF0016) |
| Potri.019G056000 | | | |
| Potri.019G057400 | AT3G10915 | | Reticulon family protein |
| Potri.019G059000 | | | |
| Potri.019G059100 | AT2G40700 | | P-loop containing nucleoside triphosphate |

| | | | |
|---|---|---|---|
| | | | hydrolases superfamily protein |
| Potri.019G059400 | AT2G40760 | | Rhodanese/Cell cycle control phosphatase superfamily protein |
| Potri.019G060900 | AT2G40770 | | zinc ion binding;DNA binding;helicases; ATP binding;nucleic acid binding |
| Potri.019G061000 | AT3G11310 | | (1 of 14) PF12776 - Myb/SANT-like DNA-binding domain (Myb_DNA-bind_3) |
| Potri.019G061400 | AT5G05190 | | Protein of unknown function (DUF3133) |
| Potri.019G068700 | | | |
| Potri.019G069300 | AT1G22880 | | cellulase 5 |
| Potri.019G088900 | | | |
| Potri.019G089100 | | | |
| Potri.019G093000 | | | |
| Potri.019G093900 | | | |
| Potri.019G094100 | | | |
| Potri.019G094700 | AT1G07560 | | Leucine-rich repeat protein kinase family protein |
| Potri.019G095700 | AT3G59690 | IQD13 | IQ-domain 13 |
| Potri.019G096500 | AT2G43780 | | |
| Potri.019G099100 | AT5G10460 | | Haloacid dehalogenase-like hydrolase (HAD) superfamily protein |
| Potri.019G102600 | | | |
| Potri.019G108700 | | | |
| Potri.019G108800 | AT4G03320 | Tic20-IV | translocon at the inner envelope membrane of chloroplasts 20-IV |
| Potri.019G112000 | AT2G31730 | | basic helix-loop-helix (bHLH) DNA-binding superfamily protein |
| Potri.019G119300 | AT4G03260 | | Outer arm dynein light chain 1 protein |
| Potri.019G119400 | | | |
| Potri.019G119700 | | | |
| Potri.019G120200 | | | |
| Potri.019G120400 | | | |
| Potri.T010600 | AT5G63620 | | GroES-like zinc-binding alcohol dehydro-genase family protein |
| Potri.T010700 | AT5G63640 | | ENTH/VHS/GAT family protein |
| Potri.T011000 | AT5G08590 | SNRK2.1 | SNF1-related protein kinase 2.1 |
| Potri.T011200 | AT5G63660 | PDF2.5 | Scorpion toxin-like knottin superfamily protein |
| Potri.T011400 | | | |
| Potri.T012200 | AT5G23630 | PDR2 | phosphate deficiency response 2 |
| Potri.T012300 | AT5G23610 | | |

| | | | |
|---|---|---|---|
| Potri.T012500 | | | |
| Potri.T012900 | | | |
| Potri.T013100 | | | |
| Potri.T044400 | AT5G08450 | | (1 of 1) KOG4843 - Uncharacterized conserved protein |
| Potri.T045400 | AT4G15093 | LigB | catalytic LigB subunit of aromatic ring-opening dioxygenase family |
| Potri.T045700 | AT5G23570 | SGS3 | XS domain-containing protein / XS zinc finger domain-containing protein-related |
| Potri.T084100 | | | |
| Potri.T084300 | AT5G60900 | | receptor-like protein kinase 1 |
| Potri.T084450 | | | |
| Potri.T085200 | AT3G57930 | | (1 of 2) PTHR34055:SF4 - EXPRESSED PROTEIN |
