## Supplemental Table 6 for "Barriers to gene flow play an important role in miantaining reproductive isolation between two closely related *Populus* (Salicaceae) species"

**Table S6. Enriched gene ontology terms among CDR genes**

| Gene ontology term | $P_{FDR}$ value | Gene count |
|---|---|---|
| response to lipid | 0.007900731 | 16（408） |
| single-organism developmental process | 0.012165981 | 33（1508） |
| cellular response to lipid | 0.012165981 | 10（201） |
| developmental process | 0.012165981 | 33（1543） |
| response to endogenous stimulus | 0.012165981 | 23（907） |
| response to oxygen-containing compound | 0.012165981 | 21（787） |
| response to hormone | 0.012165981 | 22（850） |
| brassinosteroid mediated signaling pathway | 0.012165981 | 4（25） |
| steroid hormone mediated signaling pathway | 0.012165981 | 4（25） |
| response to steroid hormone | 0.012165981 | 4（25） |
| cellular response to steroid hormone stimulus | 0.012165981 | 4（25） |
| response to alcohol | 0.012165981 | 13（364） |
| cellular response to oxygen-containing compound | 0.012165981 | 12（318） |
| cellular response to brassinosteroid stimulus | 0.012165981 | 4（26） |
| response to acid chemical | 0.012269202 | 17（589） |
| anatomical structure development | 0.012269202 | 28（1281） |
| cellular response to hormone stimulus | 0.016055699 | 13（386） |
| cellular response to endogenous stimulus | 0.016504614 | 13（390） |
| single organism reproductive process | 0.016504614 | 20（795） |
| single-multicellular organism process | 0.016504614 | 28（1325） |
| multicellular organismal process | 0.016504614 | 30（1465） |
| cellular response to chemical stimulus | 0.016504614 | 16（563） |
| developmental process involved in reproduction | 0.018295567 | 18（694） |
| reproductive process | 0.018295567 | 21（881） |
| cellular response to organic substance | 0.018295567 | 14（463） |
| reproduction | 0.018295567 | 21（883） |
| gibberellic acid mediated signaling pathway | 0.018316153 | 4（34） |
| hormone-mediated signaling pathway | 0.019095025 | 12（362） |
| gibberellin mediated signaling pathway | 0.019095025 | 4（35） |
| cellular response to gibberellin stimulus | 0.020595125 | 4（36） |
| reproductive structure development | 0.02274415 | 16（600） |
| reproductive system development | 0.02274415 | 16（600） |
| flower development | 0.025029455 | 9（228） |
| response to organic substance | 0.025029455 | 23（1056） |
| reproductive shoot system development | 0.027676478 | 9（233） |
| cellular response to acid chemical | 0.038274097 | 9（245） |
| response to stimulus | 0.039256102 | 52（3330） |
| response to abiotic stimulus | 0.040416687 | 22（1065） |
| response to brassinosteroid | 0.040416687 | 4（46） |

| | | |
|---|---|---|
| gibberellin metabolic process | 0.043435093 | 3（22） |
| protein dephosphorylation | 0.050150983 | 6（121） |

Enriched terms are displayed in color coding to reflect the relevance of ontology or functional proximity. Red, reproduction related; green, alcohol, acid chemicals related (adapted growth environment). Gene counts show the number of genes in the CDRs relative to the total number of annotated genes (in parentheses) for each term.
