## Supplemental Table 7 for "Barriers to gene flow play an important role in miantaining reproductive isolation between two closely related *Populus* (Salicaceae) species"

**Table S7.** List of the genes related to reproduction and adaptation located in regions displaying significantly high genetic differentiation between *P. alba* and *P. adenopoda*.

| Poplar gene | Best Arabidopsis hit | Synonyms | Annotated description |
|---|---|---|---|
| Potri.004G133500 | AT3G48750 | CDC2 | cell division control 2 |
| Potri.008G206000 | AT1G48270 | GCR1 | G-protein-coupled receptor 1 |
| Potri.008G217300 | AT4G02570 | CUL1 | cullin 1 |
| Potri.010G188700 | AT5G05660 | NFXL2 | sequence-specific DNA binding transcription factors; zinc ion binding; sequence-specific DNA binding transcription factors |
| Potri.010G250800 | AT3G21180 | ACA9 | autoinhibited Ca(2+)-ATPase 9 |
| Potri.012G095800 | AT5G63420 | emb2746 | RNA-metabolising metallo-beta-lactamase family protein |
| Potri.012G132400 | AT4G25420 | GA20OX1 | 2-oxoglutarate (2OG) and Fe(II)-dependent oxygenase superfamily protein |
| Potri.014G066700 | AT2G45190 | AFO | Plant-specific transcription factor YABBY family protein |
| Potri.014G164700 | AT5G35750 | HK2 | histidine kinase 2 |
| Potri.015G043200 | AT1G17980 | PAPS1 | poly(A) polymerase 1 |
| Potri.019G044400 | AT5G17690 | TFL2 | like heterochromatin protein (LHP1) |
| Potri.001G009600 | AT1G69770 | CMT3 | chromomethylase 3 |
| Potri.002G161700 | AT4G00310 | EDA8 | Putative membrane lipoprotein |
| Potri.012G100400 | AT5G63510 | MLE2.14 | gamma carbonic anhydrase like 1 |
| Potri.013G086600 | AT5G16490 | RIC4 | ROP-interactive CRIB motif-containing protein 4 |
| Potri.001G092500 | AT5G53160 | RCAR3 | regulatory components of ABA receptor 3 |
| Potri.001G215700 | AT1G08420 | BSL2 | BRI1 suppressor 1 (BSU1)-like 2 |
| Potri.002G206100 | AT2G47770 | TSPO | TSPO(outer membrane tryptophan-rich sensory protein)-related |
| Potri.002G213300 | AT4G03080 | BSL1 | BRI1 suppressor 1 (BSU1)-like 1 |

| | | | |
|---|---|---|---|
| Potri.002G233500 | AT1G10840 | TIF3H1 | translation initiation factor 3 subunit H1 |
| Potri.005G153200 | AT1G09700 | HYL1 | dsRNA-binding domain-like superfamily protein |
| Potri.005G153800 | AT4G24520 | ATR1 | P450 reductase 1 |
| Potri.007G063600 | AT4G35890 | LARP1c | winged-helix DNA-binding transcription factor family protein |
| Potri.010G164500 | AT1G14000 | VIK | VH1-interacting kinase |
| Potri.011G003400 | AT4G04740 | CPK23 | calcium-dependent protein kinase 23 |
| Potri.001G099400 | AT4G11280 | ACS6 | 1-aminocyclopropane-1-carboxy lic acid (acc) synthase 6 |
| Potri.003G064100 | AT1G17440 | EER4 | Transcription initiation factor TFIID subunit A |
| Potri.010G029500 | AT5G19000 | BPM1 | BTB-POZ and MATH domain 1 |
