## Supplemental Table 8 for "Barriers to gene flow play an important role in miantaining reproductive isolation between two closely related *Populus* (Salicaceae) species"

**Table S8.** Functional gene categories enriched for genes located within regions dis
significantly high genetic differentiation between *P. alba* and *P.adenopoda*.

| GO Term | Ontology | Description | *P*-value | FDR |
|---|---|---|---|---|
| GO:0044702 | P | single organism reproductive process | 4.90E-07 | 1.60E-03 |
| GO:0044767 | P | single-organism developmental process | 6.50E-07 | 1.60E-03 |
| GO:0032502 | P | developmental process | 1.10E-06 | 1.60E-03 |
| GO:0048856 | P | anatomical structure development | 1.20E-06 | 1.60E-03 |
| GO:0009908 | P | flower development | 1.60E-06 | 1.70E-03 |
| GO:0090567 | P | reproductive shoot system development | 2.20E-06 | 1.83E-03 |
| GO:0003006 | P | developmental process involved in reproduction | 2.40E-06 | 1.83E-03 |
| GO:0033993 | P | response to lipid | 3.00E-06 | 2.00E-03 |
| GO:0009719 | P | response to endogenous stimulus | 3.60E-06 | 2.13E-03 |
| GO:0022414 | P | reproductive process | 6.20E-06 | 3.11E-03 |
| GO:0048608 | P | reproductive structure development | 7.00E-06 | 3.11E-03 |
| GO:0061458 | P | reproductive system development | 7.00E-06 | 3.11E-03 |
| GO:0009725 | P | response to hormone | 9.90E-06 | 3.80E-03 |
| GO:0044707 | P | single-multicellular organism process | 1.00E-05 | 3.80E-03 |
| GO:1901700 | P | response to oxygen-containing compound | 1.10E-05 | 3.91E-03 |
| GO:0009909 | P | regulation of flower development | 1.30E-05 | 4.33E-03 |
| GO:0032501 | P | multicellular organismal process | 1.50E-05 | 4.70E-03 |
| GO:0007275 | P | multicellular organismal development | 3.30E-05 | 9.76E-03 |
| GO:0001101 | P | response to acid chemical | 3.60E-05 | 1.01E-02 |
| GO:0048731 | P | system development | 4.00E-05 | 1.07E-02 |
| GO:0097305 | P | response to alcohol | 4.90E-05 | 1.24E-02 |
| GO:0010033 | P | response to organic substance | 5.90E-05 | 1.43E-02 |
| GO:2000241 | P | regulation of reproductive process | 6.30E-05 | 1.46E-02 |
| GO:0048831 | P | regulation of shoot system development | 8.50E-05 | 1.85E-02 |
| GO:0048367 | P | shoot system development | 8.70E-05 | 1.85E-02 |
| GO:0009791 | P | post-embryonic development | 0.00011 | 2.25E-02 |
| GO:0071396 | P | cellular response to lipid | 0.00015 | 2.96E-02 |
| GO:0009742 | P | brassinosteroid mediated signaling pathway | 2.00E-04 | 3.55E-02 |
| GO:0043401 | P | steroid hormone mediated signaling pathway | 2.00E-04 | 3.55E-02 |
| GO:0071383 | P | cellular response to steroid hormone stimulus | 2.00E-04 | 3.55E-02 |
| GO:0071367 | P | cellular response to brassinosteroid stimulus | 0.00022 | 3.78E-02 |
| GO:0048545 | P | response to steroid hormone | 0.00025 | 4.16E-02 |
| GO:0009628 | P | response to abiotic stimulus | 0.00029 | 4.68E-02 |
| GO:0032870 | P | cellular response to hormone stimulus | 4.00E-04 | 6.27E-02 |
| GO:0071495 | P | cellular response to endogenous stimulus | 0.00043 | 6.54E-02 |
| GO:1901701 | P | cellular response to oxygen-containing compound | 5.00E-04 | 7.34E-02 |

| | | | | |
|---|---|---|---|---|
| GO:0048580 | P | regulation of post-embryonic development | 0.00051 | 7.34E-02 |
| GO:0009755 | P | hormone-mediated signaling pathway | 0.00066 | 9.25E-02 |
| GO:0030154 | P | cell differentiation | 0.00069 | 9.42E-02 |
| GO:0009740 | P | gibberellic acid mediated signaling pathway | 0.00077 | 1.03E-01 |
| GO:0071310 | P | cellular response to organic substance | 0.00083 | 1.05E-01 |
| GO:0070887 | P | cellular response to chemical stimulus | 0.00083 | 1.05E-01 |
| GO:0010476 | P | gibberellin mediated signaling pathway | 0.00091 | 1.13E-01 |
| GO:0071370 | P | cellular response to gibberellin stimulus | 0.00099 | 1.20E-01 |
| GO:0048869 | P | cellular developmental process | 0.00143 | 1.69E-01 |
| GO:0051239 | P | regulation of multicellular organismal process | 0.00148 | 1.71E-01 |
| GO:0009685 | P | gibberellin metabolic process | 0.00162 | 1.84E-01 |
| GO:0009739 | P | response to gibberellin | 0.00169 | 1.88E-01 |
| GO:0065007 | P | biological regulation | 0.00174 | 1.89E-01 |
| GO:0007015 | P | actin filament organization | 0.00178 | 1.90E-01 |
| GO:0050789 | P | regulation of biological process | 0.00209 | 2.14E-01 |
| GO:2000026 | P | regulation of multicellular organismal development | 0.00215 | 2.14E-01 |
| GO:0009737 | P | response to abscisic acid | 0.00216 | 2.14E-01 |
| GO:0030036 | P | actin cytoskeleton organization | 0.00217 | 2.14E-01 |
| GO:0050896 | P | response to stimulus | 0.00228 | 2.15E-01 |
| GO:0030029 | P | actin filament-based process | 0.00231 | 2.15E-01 |
| GO:1902182 | P | shoot apical meristem development | 0.00237 | 2.15E-01 |
| GO:1902183 | P | regulation of shoot apical meristem development | 0.00237 | 2.15E-01 |
| GO:0050794 | P | regulation of cellular process | 0.00238 | 2.15E-01 |
| GO:0009741 | P | response to brassinosteroid | 0.00261 | 2.32E-01 |
| GO:0009416 | P | response to light stimulus | 0.0027 | 2.36E-01 |
| GO:0007389 | P | pattern specification process | 0.00289 | 2.48E-01 |
| GO:0009888 | P | tissue development | 0.00299 | 2.50E-01 |
| GO:0009798 | P | axis specification | 0.00303 | 2.50E-01 |
| GO:0071229 | P | cellular response to acid chemical | 0.00305 | 2.50E-01 |
| GO:0042221 | P | response to chemical | 0.00311 | 2.51E-01 |
| GO:0050829 | P | defense response to Gram-negative bacterium | 0.00377 | 2.98E-01 |
| GO:0097306 | P | cellular response to alcohol | 0.0038 | 2.98E-01 |
| GO:0050793 | P | regulation of developmental process | 0.00411 | 3.17E-01 |
| GO:0009314 | P | response to radiation | 0.0045 | 3.39E-01 |
| GO:0006075 | P | (1->3)-beta-D-glucan biosynthetic process | 0.00458 | 3.39E-01 |
| GO:0010048 | P | vernalization response | 0.00458 | 3.39E-01 |
| GO:0009793 | P | embryo development ending in seed dormancy | 0.00471 | 3.39E-01 |
| GO:0009733 | P | response to auxin | 0.00474 | 3.39E-01 |
| GO:0006470 | P | protein dephosphorylation | 0.00477 | 3.39E-01 |
| GO:0071407 | P | cellular response to organic cyclic compound | 0.00519 | 3.63E-01 |
| GO:0044699 | P | single-organism process | 0.00532 | 3.63E-01 |

| | | | | |
|---|---|---|---|---|
| GO:0016101 | P | diterpenoid metabolic process | 0.00543 | 3.63E-01 |
| GO:0006074 | P | (1->3)-beta-D-glucan metabolic process | 0.00546 | 3.63E-01 |
| GO:0010310 | P | regulation of hydrogen peroxide metabolic process | 0.00546 | 3.63E-01 |
| GO:0014070 | P | response to organic cyclic compound | 0.0068 | 4.47E-01 |
| GO:0009790 | P | embryo development | 0.00723 | 4.70E-01 |
| GO:0090056 | P | regulation of chlorophyll metabolic process | 0.00744 | 4.71E-01 |
| GO:0048509 | P | regulation of meristem development | 0.0075 | 4.71E-01 |
| GO:0003002 | P | regionalization | 0.00752 | 4.71E-01 |
| GO:0048507 | P | meristem development | 0.0076 | 4.71E-01 |
| GO:0009735 | P | response to cytokinin | 0.008 | 4.76E-01 |
| GO:0006188 | P | IMP biosynthetic process | 0.00853 | 4.76E-01 |
| GO:0008356 | P | asymmetric cell division | 0.00968 | 4.76E-01 |
| GO:0046040 | P | IMP metabolic process | 0.00968 | 4.76E-01 |
| GO:1901401 | P | regulation of tetrapyrrole metabolic process | 0.00968 | 4.76E-01 |
| GO:0009266 | P | response to temperature stimulus | 0.00974 | 4.76E-01 |
| GO:0010629 | P | negative regulation of gene expression | 0.00993 | 4.80E-01 |
| GO:0009409 | P | response to cold | 0.01 | 4.80E-01 |
| GO:0000148 | C | 1,3-beta-D-glucan synthase complex | 0.0046 | 1.00E+00 |
| GO:0044464 | C | cell part | 0.0078 | 1.00E+00 |
| GO:0004652 | F | polynucleotide adenylyltransferase activity | 0.00047 | 4.52E-01 |
| GO:0045544 | F | gibberellin 20-oxidase activity | 0.00047 | 4.52E-01 |
| GO:0003700 | F | transcription factor activity, sequence-specific DNA binding | 0.00069 | 4.52E-01 |
| GO:0001071 | F | nucleic acid binding transcription factor activity | 7.00E-04 | 4.52E-01 |
| GO:0004449 | F | isocitrate dehydrogenase (NAD+) activity | 0.00116 | 5.30E-01 |
| GO:0048029 | F | monosaccharide binding | 0.00123 | 5.30E-01 |
| GO:0048027 | F | mRNA 5'-UTR binding | 0.00161 | 5.95E-01 |
| GO:0016706 | F | oxidoreductase activity, acting on paired donors, wi incorporation or reduction of molecular oxygen, 2-oxoglutarate as one donor, and incorporation of one atom each of oxygen into both donors | 0.00237 | 7.66E-01 |
| GO:0001085 | F | RNA polymerase II transcription factor binding | 0.004 | 1.00E+00 |
| GO:0003843 | F | 1,3-beta-D-glucan synthase activity | 0.00412 | 1.00E+00 |
| GO:0004448 | F | isocitrate dehydrogenase activity | 0.00491 | 1.00E+00 |
| GO:0008134 | F | transcription factor binding | 0.00521 | 1.00E+00 |
| GO:0000977 | F | RNA polymerase II regulatory region sequence-sp DNA binding | 0.0057 | 1.00E+00 |
| GO:0001012 | F | RNA polymerase II regulatory region DNA bindin | 0.0057 | 1.00E+00 |
| GO:0004525 | F | ribonuclease III activity | 0.00767 | 1.00E+00 |
| GO:0032296 | F | double-stranded RNA-specific ribonuclease activit | 0.00767 | 1.00E+00 |
