## Supplemental Table 9 for "Barriers to gene flow play an important role in miantaining reproductive isolation between two closely related *Populus* (Salicaceae) species"

**Table S9.** Gene introduction

1. **Cell division.** The CDC2 (cyclin-dependent kinase) gene encodes a catalytic subunit of a cyclin-dependent kinase and is involved in many cellular processes, including the control of cell division. Many of the cdc2 gene and its related genes encode proteins that interact with cyclins; these genes are collectively referred to as cyclin-dependent kinases (Rossmacdonald et al., 1994). Fobert et al. isolated four cdc2-related genes from Antirrhinum (Fobert et al., 1996). Genetic studies of cell division in yeast have identified the product of the cdc2 gene as a key component of these cascades. And the histological distribution of their transcripts suggests that members of the Antirrhinum cdc2 gene family are differentially expressed during the cell division cycle (Fobert et al., 1996).

2. **Pollen rejection pathways.** Mutations in CUL1 (Cullin1) establish unilateral incompatibility with self-incompatibility (SI) populations and strengthen reproductive isolation (Markova et al., 2016). Li et al. used a self-incompatible wild tomato species Solanum Arcanum, silence CUL1 expression in pollen by RNA interference (RNAi), resulting in pollen rejection in normally compatible sibling hybridization, while the same pollen remained fully compatible on sc logins expressing inactive s-RNase. The results strongly suggest that CUL1 has a role in protecting pollen from s-rnases in SI and UI (interspecific incompatibility) and provides further evidence for overlap between intraspecific and interspecific pollen rejection pathways (Li & Chetelat, 2014).

3. **Embryogenesis.** Emb2746 genes play an important role in fruit development. In the GT-1 subfamily, EMB2746 (At5g63420), which exists in *Arabidopsis thaliana*, is widely expressed in the vegetative part of Arabidopsis thaliana, especially in seeds (Ma et al., 2019). The seeds of the emb2746 mutant only develop to the globular stage; therefore, EMB2746 is necessary for early embryogenesis. LOC_Os02g33610 in rice is also similar to At5g63420 (Qin et al., 2014).

4. **Rice flowers/spike into seedlings and reproductive habits.** Wang et al. found three naturally occurring mutants in rice, namely phoenix (PHO), degenerative paleo

(DEP), and abnormal floral organs (AFO) (Wang et al., 2010). By analyzing three naturally occurring mutants in rice, they found that mutations in DEP and AFO led to the transformation of rice flowers/spike into seedlings, and subsequently to the shift of reproductive strategies from sexual to non-sexual, suggesting that DEP and AFO may synergize to regulate rice reproductive habits (Kumaran et al., 1999; Wang et al., 2010).

5. **Plant germ cell development during meiosis.** The HK2 (Histidine kinase 2) gene has an important relationship with the gene regulatory mechanisms that are crucial to plant germ cell development during meiosis. HK2 gene regulates female meiosis by modulating meiotic gene expression (Nishimura et al., 2004).

6. **Transition from photoperiod regulation to reproductive growth.** TFL2 (TERMINAL FLOWER2) gene itself may regulate numerous developmental processes and TFL2 specifically represses FLOWERING LOCUS T (FT) in the flowering pathway (Kotake et al., 2003). Larsson et al. used *Arabidopsis thaliana* to describe a novel definitive mutant, terminal flower 2 (TFL2). Besides its role in maintaining the identity of inflorescence meristem, TFL2 is also active in the signal-aware pathway of the transition from photoperiod regulation to reproductive growth (Larsson et al., 1998).

7. **The early development of plant embryos and pollen tubes.** EDA8 (Embryo sac development) gene is a gene expressed from the female gametophyte or the maternal genome and plays an important role in the early development of plant embryos. Zhu et al. found a gene embryo sac 1 (OSEMSA1) containing lysosomal domains involved in the sexual reproduction of rice, this gene encodes a protein containing a lysosomal domain that is required for blastocyst development and function (Zhu et al., 2017). The gene is expressed in roots, stems, leaf tissues, spikes, and ovaries and plays a role in hormone regulation. Inhibition of osemsa1 expression leads to blastocyst defects and poor differentiation of gametophyte cells, which can not attract pollen tubes, thus reducing the rate of spike seed formation (Zhu et al., 2017).
